# Evolutionary Rewiring of Anthocyanin Biosynthesis Pathway in *Poaceae*

**DOI:** 10.1101/2025.09.21.677584

**Authors:** Najnin Khatun, Aidan Jones, Avery Rahe, Nancy Choudhary, Boas Pucker

## Abstract

*Poaceae* is a species-rich family of plants in the monocots harboring many important crop species. As grasses, these plants lack striking flowers that are typically pigmented by anthocyanins.

Here, we investigated the anthocyanin biosynthesis genes in Poaceae and characterized several striking patterns. (1) An independently evolved MYB lineage (ncaMYB) is responsible for the regulation of the anthocyanin biosynthesis in Poaceae. (2) The independently evolved BZ2 lineage replaced the otherwise widespread TT19/An9 lineage as anthocyanin-related glutathione S-transferase in Poaceae. (3) The anthocyanin biosynthesis genes *ANS, arGST (BZ2)*, and the anthocyanin biosynthesis activating MYB (ncaMYB) appear to be absent from the genus *Brachypodium*.

These observations could be explained by a loss of the anthocyanin biosynthesis at the basis of the *Poaceae*, followed by an evolutionarily independent regain and again a loss of the anthocyanin biosynthesis in *Brachypodium*.

## Introduction

Anthocyanins are well known as pigments conferring striking colors to flowers, but have several additional functions in stress response and plant-animal interactions (Grünig *et al*., 2024). As pigments, anthocyanins can protect plant structures from intense light exposure (Gould *et al*., 1995; Feild *et al*., 2001; Grünig *et al*., 2024) and contribute to the scavenging of reactive oxygen species due to their antioxidant properties (Gould *et al*., 2002; Nagata *et al*., 2003; Kytridis & Manetas, 2006; Grünig *et al*., 2024). While protection is most likely the primary function in leaves and other vegetative plant structures, the coloration of flowers and fruits is considered an important signal to attract animals (Weiss, 1991; Willmer *et al*., 2009; Ruxton & Schaefer, 2016; Garcia *et al*., 2022; Grünig *et al*., 2024). The diversity of functions might explain why plants produce a plethora of different anthocyanin derivatives. Differences in the hydroxylation of the central B-ring result in different colors: one hydroxylation leads to orange, two hydroxylations lead to pink, and three hydroxylations result in blue (Winkel-Shirley, 2001; Bogs *et al*., 2006; Schwinn *et al*., 2014). Additional modifications with glycosyl, acyl, methyl, and other groups can alter the color and biochemical properties of anthocyanins (Grünig *et al*., 2024). Anthocyanins are produced by one branch of the flavonoid biosynthesis that competes for substrate with other branches, leading to flavonols and proanthocyanidins (PAs) (Winkel-Shirley, 2001; Yuan *et al*., 2016; Choudhary & Pucker, 2024). Activity of different branches is controlled at the transcriptional level by several transcription factors, most notably members of the MYB family (Nesi *et al*., 2001; Stracke *et al*., 2007; Gonzalez *et al*., 2008).

Transcriptional regulation of the anthocyanin biosynthesis is controlled by a complex of different transcription factors and considered a model system for gene expression in eukaryotes. A transcription factor complex, the MBW complex – named after the MYB, bHLH, and WD40 proteins forming it-is a well-known activator of anthocyanin biosynthesis genes (Gonzalez *et al*., 2008; Lloyd *et al*., 2017). MBW complexes were studied across a wide range of plant species (Ramsay & Glover, 2005; Lang *et al*., 2010; Feller *et al*., 2011; Xu *et al*., 2015; Zhang & Schrader, 2017; Zhang *et al*., 2019). The integration of different MYB and bHLH proteins leads to specificity for certain target genes, which explains the control of various processes in development and stress response beyond the anthocyanin biosynthesis (Zhang & Schrader, 2017; Zhang *et al*., 2019). The conserved WD40 proteins are hypothesized to be a scaffold to enhance gene transcription mediated by the MYB-bHLH complex (Ramsay & Glover, 2005). Multiple studies corroborated that the MYB protein is the most important factor for the specificity of the MBW complex towards anthocyanin biosynthesis genes (Zhang & Schrader, 2017; Zhang *et al*., 2019; Marin-Recinos & Pucker, 2024).

Anthocyanins are synthesized based on phenylalanine that is channeled through the general phenylpropanoid pathway and a specific branch of the flavonoid biosynthesis (**Fig. 1**). The first committed step of the flavonoid biosynthesis is the chalcone synthase (CHS) that generates naringenin chalcone (Ferrer *et al*., 1999). The conversion of naringenin chalcone into naringenin is catalyzed by the chalcone isomerase (CHI) (Jez *et al*., 2000). The flavanone 3-hydroxylase (F3H) adds a hydroxy group, resulting in dihydroflavonols (Forkmann *et al*., 1980). A competing reaction catalyzed by the flavone synthase (FNS) channels naringenin into the flavone biosynthesis (Martens & Forkmann, 1999; Martens *et al*., 2003). Dihydroflavonols are the substrate of dihydroflavonol 4-reductase (DFR) and flavonol synthase (FLS) that form the branching point between anthocyanin biosynthesis and flavonol biosynthesis (Davies *et al*., 2003; Choudhary & Pucker, 2024). The next step in the anthocyanin biosynthesis is the conversion of leucoanthocyanidins into anthocyanidins by the anthocyanidin synthase (ANS). This process also requires the anthocyanin-related glutathione-S-transferase (arGST) (Eichenberger *et al*., 2023). Finally, the addition of a sugar moiety and further additions of various groups lead to the formation of diverse anthocyanins (Tohge *et al*., 2005; Luo *et al*., 2007; Yonekura-Sakakibara *et al*., 2012; Grünig *et al*., 2024).

**Fig. 1:**
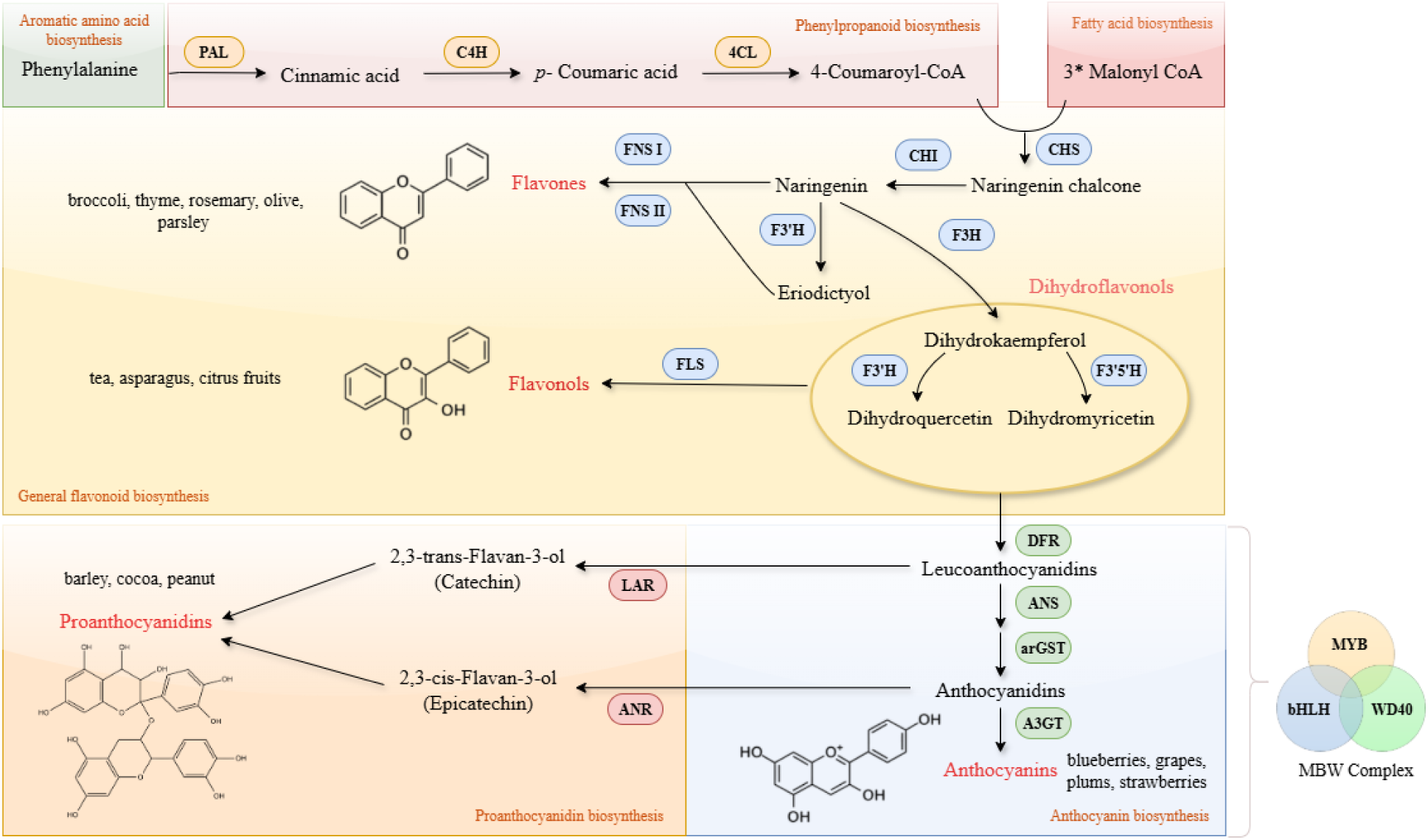
Simplified illustration of the flavonoid biosynthesis. The enzyme abbreviations are as follows: PAL: Phenylalanine ammonia-lyase, C4H: Cinnamic acid 4-hydroxylase, 4CL: 4-coumarate-CoA ligase, CHS: Chalcone synthase, CHI: Chalcone isomerase, F3H: Flavanone 3-hydroxylase, F3’H: Flavonoid 3’-hydroxylase, F3’5’H: Flavonoid 3’,5’-hydroxylase, FNS: Flavone synthase. FLS: Flavonol synthase, DFR: Dihydroflavonol 4-reductase, ANS: Anthocyanidin synthase, arGST: Anthocyanin-related glutathione S-transferase, A3GT: Anthocyanidin 3-O-glycosyl transferase, ANR: Anthocyanidin reductase, LAR: Leucoanthocyanidin reductase, MBW complex: MYB (v-myb avian myeloblastosis viral oncogene homolog), bHLH (basic helix-loop-helix), and WD40.

Branching from the anthocyanin biosynthesis is the proanthocyanidin biosynthesis. Leucoanthocyanidin reductase (LAR) catalyzes the conversion of leucocyanidin to (+)-catechin (Tanner *et al*., 2003). More recent evidence also shows that LAR also plays a role in the polymerization of proanthocyanidin subunits in some species by controlling starter unit availability (Liu *et al*., 2016).

Anthocyanin biosynthesis takes place in the cytosol, specifically at the endoplasmic reticulum (ER) (Pucker & Selmar, 2022; Ma *et al*., 2024). An ER-bound metabolon comprising enzymes of the anthocyanin biosynthesis has been proposed and supported by several protein-protein interaction studies (Crosby *et al*., 2011; Fujino *et al*., 2018; Nakayama *et al*., 2019; Ruan *et al*., 2024). After synthesis in the cytosol, anthocyanins accumulate in the central vacuole of plant cells (Zhao & Dixon, 2010; Pucker & Selmar, 2022). It has been proposed that an anthocyanin-related glutathione S-transferase (arGST) acts as ligandin during the transport of anthocyanins into the central vacuole (Grotewold, 2006; Petrussa *et al*., 2013; Pucker & Selmar, 2022). Two alternative theories have been postulated (i) the vesicle trafficking model and (ii) the transporter model (Grotewold & Davies, 2008; Zhao, 2015; Pucker & Selmar, 2022). Both models propose that a glutathione S-transferase (GST) plays a key role in the efficient transport of anthocyanins from the endoplasmic reticulum (ER) to the central vacuole of the plant (Grotewold & Davies, 2008; Petrussa *et al*., 2013; Pucker & Selmar, 2022). Not all GSTs are associated with the anthocyanin biosynthesis. GST proteins have been classified into 14 different classes, including theta (T), tau (U), phi (F), zeta (Z), lambda (L), and iota (I) (Liu *et al*., 2013; Lallement *et al*., 2014; Hernández Estévez & Rodríguez Hernández, 2020). Among them, tau (U) and phi (F) are the two most abundant classes in land plants (Liu *et al*., 2013; Lallement *et al*., 2014; Hernández Estévez & Rodríguez Hernández, 2020).

The involvement of GST in anthocyanin accumulation was initially established in *Zea mays* through the study of bronze-2 (bz2) mutant (Marrs *et al*., 1995). Numerous arGST proteins have been identified across diverse plant species, including *TT19* in *Arabidopsis thaliana* (Kitamura *et al*., 2004; Sun *et al*., 2012), *RsGST1* in *Raphanus sativus* (Lai *et al*., 2021), *GST1* in *Perilla frutescens* (Yamazaki *et al*., 2008), *VvGST1* and *VvGST4* in *Vitis vinifera* (Conn *et al*., 2008; Pérez-Díaz *et al*., 2016), *AcGST1* in *Actinidia chinensis* (Liu *et al*., 2019b), *LcGST4* in *Litchi chinensis* Sonn (Hu *et al*., 2016). The ability of arGSTs from different plant species to complement mutants in distantly related plant lineages was explored (Alfenito *et al*., 1998). For instance, the introduction of *Zea mays arGST* (*ZmBZ2*) cDNA by particle bombardment method is capable of complementing mutations in the corresponding genes encoding *arGST* (*PhAN9*) in *Petunia* sp. and vice versa (Alfenito *et al*., 1998). *ZmBZ2* and *PhAN9* are capable of complementing an arGST mutation in carnation (Larsen *et al*., 2003).

While originally an enzymatic function of arGST in the anthocyanin metabolism was proposed (Marrs *et al*., 1995; Alfenito *et al*., 1998), lack of evidence for an enzymatic activity led to the postulation of a role as ligandin in the anthocyanin transport (Mueller *et al*., 2000). A recent study provides evidence that arGSTs have enzymatic functions in the anthocyanin biosynthesis beyond intracellular transport and even questions a role in anthocyanin transport (Eichenberger *et al*., 2023). Additionally, epicatechin, a key precursor of proanthocyanidins, is cytosolically produced from cyanidin by the enzyme anthocyanidin reductase (ANR) (Lu *et al*., 2022). Notably, in *Arabidopsis thaliana,* the *tt19 arGST* mutation eliminates the epicatechin production completely (Kitamura *et al*., 2010). This outcome is unexpected in the frame of the ligandin model, as mutating a transporter protein should not affect the cytosolic availability of cyanidin, the precursor of the epicatechin.

While arGSTs are now known to play important roles in anthocyanin biosynthesis (Eichenberger *et al*., 2023), it remains unclear whether orthologs from various plant species behave similarly or exhibit functional divergence. In investigating the functional divergence of arGSTs and other structural genes in the anthocyanin biosynthesis pathway across plant lineages, we have focused on *Poaceae*, considering its ecological and agricultural importance as well as the importance of maize for early research on pigmentation (McClintock, 1950; Cone *et al*., 1986; Marrs *et al*., 1995). As the fourth-largest plant family, *Poaceae* consists of over 12,000 species, including integral cereal crops such as *Avena sativa*, *Hordeum vulgare*, *Triticum aestivum*, *Oryza sativa*, *Sorghum bicolor*, and *Zea mays* (Soreng *et al*., 2022). Phylogenetically, *Poaceae* is divided into two major clades, the BOP clade (*Bambusoideae*, *Oryzoideae*, and *Pooideae*) and the PACMAD clade (*Panicoideae, Aristidoideae, Chloridoideae, Micrairoideae, Arundinoideae, and Danthonioideae*). Following their divergence at the base of the core *Poaceae*, these lineages have gained distinct physiological and ecological adaptations, as reflected through characteristic differences such as in life histories, habitat, morphology, average genome sizes, and predominant photosynthetic pathways (Taylor *et al*., 2010; Simpson *et al*., 2024).

The flavonoid biosynthesis is well known for numerous gene duplications (Choudhary & Pucker, 2024; Marin-Recinos & Pucker, 2025), which can result in sub– and neofunctionalization and contribute to the evolution of the complexity seen today (Falginella *et al*., 2010; Kawai *et al*., 2014; Katsu *et al*., 2017; Jia *et al*., 2019; Pucker & Iorizzo, 2023). Many plant species harbour several gene copies for the different steps of the flavonoid biosynthesis, often connected to tissue-specific expression of these copies (Falginella *et al*., 2010; Katsu *et al*., 2017; Schilbert *et al*., 2021). However, there is also the opposite pattern: loss of a gene function – frequently seen for the anthocyanin and proanthocyanidin biosynthesis branches due to the visible phenotypes (Appelhagen *et al*., 2014; Marin-Recinos & Pucker, 2024; Horz *et al*., 2024). While anthocyanin loss occurs frequently throughout all major plant lineages, it usually affects only individual species or accessions within a species (Onozaki *et al*., 1999; Zufall & Rausher, 2003; Smith & Rausher, 2011; Marin-Recinos & Pucker, 2024). Most known cases can be attributed to changes in transcription regulation, e.g. loss of the anthocyanin-specific MYB transcription factor (Marin-Recinos & Pucker, 2024). Loss at a higher taxonomic level has only been reported for several families in the Caryophyllales, where the anthocyanin pigmentation has been replaced by betalain pigmentation (Timoneda *et al*., 2019). Here, loss of the transcriptional regulation by the MBW complex and wholesome loss of the arGST gene have been identified as important factors (Pucker *et al*., 2024b). Several studies have reported the absence of proanthocyanidin biosynthesis genes at the species level, e.g. absence of *LAR* in *Zea mays* and *Sorghum bicolor* (Lu *et al*., 2023). Despite this loss, however, *Zea mays* is still able to synthesize catechin-cysteine from 2,3-*trans*-leucocyanidin through an unknown mechanism (Lu *et al*., 2023). Along with LAR, anthocyanidin reductase (ANR) is an integral enzyme in proanthocyanidin synthesis, converting cyanidins to flavan-3-ol starter and extension subunits required for proanthocyanidin polymerization (Xie *et al*., 2003; Jun *et al*., 2021). While ANR in eudicot species commonly produces (-)-epicatechin, the ANR homolog in *Zea mays* preferentially produces (+)-epicatechin and 4β-(S-cysteinyl)-catechin, giving rise to unusual procyanidin dimers (Jun *et al*., 2021; Lu *et al*., 2023). A major loss event of the proanthocyanidin biosynthesis gene *LAR* was recently reported in the *Brassicaceae* (Marin-Recinos & Pucker, 2025).

Here, we investigated the evolution of the anthocyanin biosynthesis with a particular focus on *Poaceae*. Our results suggest (1) an independent MYB lineage for anthocyanin activation in the *Poaceae*, (2) an independent evolution of arGST in *Poaceae*, (3) complex loss/gain of proanthocyanidin biosynthesis genes in *Poaceae*, and (4) the loss of the anthocyanin biosynthesis in *Brachypodium*.

## Materials & Methods

### Data collection

Numerous plant genome sequences representing diverse taxonomic groups were collected as the basis for gene expression analysis using public RNA-seq data (Pucker *et al*., 2024a, 2025). To enable a focused investigation of anthocyanin biosynthesis in *Poaceae*, additional *Poaceae* datasets and datasets from other monocots and dicots were included as outgroups (**Table S1 in Additional File 1**). Homology-based gene prediction of 13 *Poaceae* genome sequences lacking annotation was performed using GeMoMa v1.9 (Keilwagen *et al*., 2016, 2019) based on existing high-quality annotations of eight *Poaceae* genomes (*Triticum turgidum* (Maccaferri *et al*., 2019), *Alopecurus aequalis* (Wright *et al*., 2024), *Brachypodium distachyon* (Vogel *et al*., 2010), *Aegilops tauschii* (Luo *et al*., 2017), *Triticum aestivum* (Zhu *et al*., 2021), *Lolium rigidum* (Paril *et al*., 2022), *Hordeum vulgare* (Beier *et al*., 2017), and *Triticum urartu* (Ling *et al*., 2018)). For each of the eight reference species, extracted coding sequences were aligned to the sequences of the 13 target genomes using MMSeqs2 (version 17-b804f)(Steinegger & Söding, 2017), with default parameters applied by GeMoMa. The resulting eight gene annotation sets were filtered and merged using GeMoMa Annotation Filter (GAF) with parameters f = “start = =‘M’ and stop = =‘*’ and (isNaN(score) or score/aa > =1.52)” and atf=“sumWeight>4” for all annotated species. Assembly and annotation completeness were assessed with BUSCO v5.8.2 (Manni *et al*., 2021) using the *embryophyta_odb12* lineage dataset (**Table S2 in Additional File 1**).

### Flavonoid biosynthesis gene identification

KIPEs v3 (Rempel *et al*., 2023) was used to identify the functional structural genes of the anthocyanin biosynthesis pathway, including *F3H*, *FLS*, *DFR*, *ANS*, *arGST*, *LAR*, and *ANR,* in all these datasets.

KIPEs v3 provided the required bait sequences and essential parameters, facilitating the accurate identification of our target genes. To identify MYB sequences, anthocyanin biosynthesis activator MYBs described in the literature were collected. Among these, monocot and Arabidopsis MYBs were used as baits to identify similar sequences in all datasets using the python script collect_best_BLAST_hits.py available at https://github.com/bpucker/ApiaceaeFNS1 (Pucker & Iorizzo, 2023).

### Co-expression analysis

RNA-seq data sets were processed with kallisto v0.44 (Bray *et al*., 2016), utilizing the coding sequences of each plant species. The Python script kallisto_pipeline3.py (Pucker & Iorizzo, 2023) was employed to facilitate the execution of kallisto across all input files. Subsequently, the merge_kallisto_output3.py script (Pucker & Iorizzo, 2023) was used to merge the individual kallisto output files into a count table for each species and gene count normalization by the transcripts per million (TPM). The transcript abundance data were then used in the subsequent co-expression analysis with CoExp (https://github.com/bpucker/CoExp) to validate the accurate identification of arGST sequences.

Functional annotations for all plant species were generated using *Arabidopsis thaliana* as a model system. A Python script construct_anno.py was used to annotate all plant species (https://github.com/bpucker/PlantGenomicsGuide).

### Phylogenetic analysis

Phylogenetic trees were constructed for anthocyanin-related MYB sequences, the 2ODD family (*F3H*, *FLS*, and *ANS*), the SDR superfamily (*LAR*, *ANR*, and *DFR*), and the arGST family. For the MYB tree, sequences showing at least 40% similarity to the bait sequences were used as input. For structural genes, sequences with more than 70% residue conservation in KIPEs were included. Previously reported functionally characterized sequences were also incorporated into all trees. For the arGST tree, additional GST sequences, such as Tau, Phi, Lambda, Zeta, and their orthologs, were retrieved from NCBI (Sayers *et al*., 2025) and TAIR (Lamesch *et al*., 2012) and used as outgroups for tree construction. Global alignments of all candidate sequences were generated with MAFFT v7.310 (Katoh & Standley, 2013) using the G-INS-i method with default gap penalties. The resulting amino acid alignments were back-translated into codon alignments with pxaa2cdn from Phyx (Brown *et al*., 2017). The gaps within the alignments were subsequently refined using trimAl (--automated 1) (Capella-Gutiérrez *et al*., 2009). The phylogenetic trees were inferred with IQ-TREE v2.3.6 (Nguyen *et al*., 2015; Minh *et al*., 2020) using 1000 bootstrap replicates for statistical support. The best-fit substitution model was determined using ModelFinder (Kalyaanamoorthy *et al*., 2017). Additional phylogenetic trees were constructed based on MAFFT v7.310 (Katoh & Standley, 2013) and Muscle v5.3 (Edgar, 2022) with IQ-TREE v2.3.6 (Minh *et al*., 2020) and FastTree v2.2.0 (Price *et al*., 2010). The topologies of the resulting trees were manually compared to validate the crucial nodes supporting major conclusions (**Fig. S1-S14 in Additional File 1**). Phylogenetic trees were visualized in iTOL v7 (Letunic & Bork, 2024).

### Microsynteny analysis

JCVI/MCScan v1.5.6 (Tang *et al*., 2015) was used to compare local synteny among the genome sequences of Brachypodium species and outgroups belonging to the *Poales* order. The species *Ananas comosus*, *Zea mays*, *Aegilops tauschii*, *Hordeum marinum*, *Brachypodium arbuscula*, *Brachypodium distachyon*, *Brachypodium hybridum*, *Brachypodium mexicanum*, *Brachypodium stacei*, and *Brachypodium sylvaticum* were included. The regions surrounding MYB C1/Pl and ANS were selected to investigate the fate of these genes in the *Poaceae* family.

## Results and Discussions

### 1. Independent evolution of *Poaceae* anthocyanin activator MYB

The identity of the constituent MYB largely determines the specific function of the MBW complex. In the proanthocyanidin pathway, *AtTT2* and its homologs in other species are known to determine PAs accumulation in seeds and are called TT2/PA2 MYBs (Nesi *et al*., 2001; Terrier *et al*., 2009).

Interestingly, in some plants, another type of MYB evolved in PA accumulation, called the PA1-type (Bogs *et al*., 2007; Ravaglia *et al*., 2013; Wang *et al*., 2018). This PA1 MYB is not an ortholog of AtTT2 and appears in a different lineage (**Fig. 2**). The functionally redundant *Arabidopsis* MYB transcription factors MYB75 (PAP1), MYB90 (PAP2), MYB113, and MYB114 are known to activate anthocyanin biosynthesis pathway genes (Gonzalez *et al*., 2008; Li, 2014). However, *Zea mays* has the tissue-specific anthocyanin biosynthesis activators C1 and Pl (Cone *et al*., 1993). OsC1 in *Oryza sativa* is a homolog of ZmC1/ZmPl (Sakamoto *et al*., 2001). Our phylogenetic analysis reveals that *Poaceae* anthocyanin MYB sequences do not cluster with canonical anthocyanin MYB (caMYB), but rather with Pl and C1 genes from *Zea mays* and *Oryza sativa,* hereafter called the non-canonical anthocyanin MYBs (ncaMYB) (**Fig. 2**). *Brachypodium* sequences do not have a ncaMYB ortholog and are absent in the *Poaceae*-specific ncaMYB clade.

**Fig. 2:**
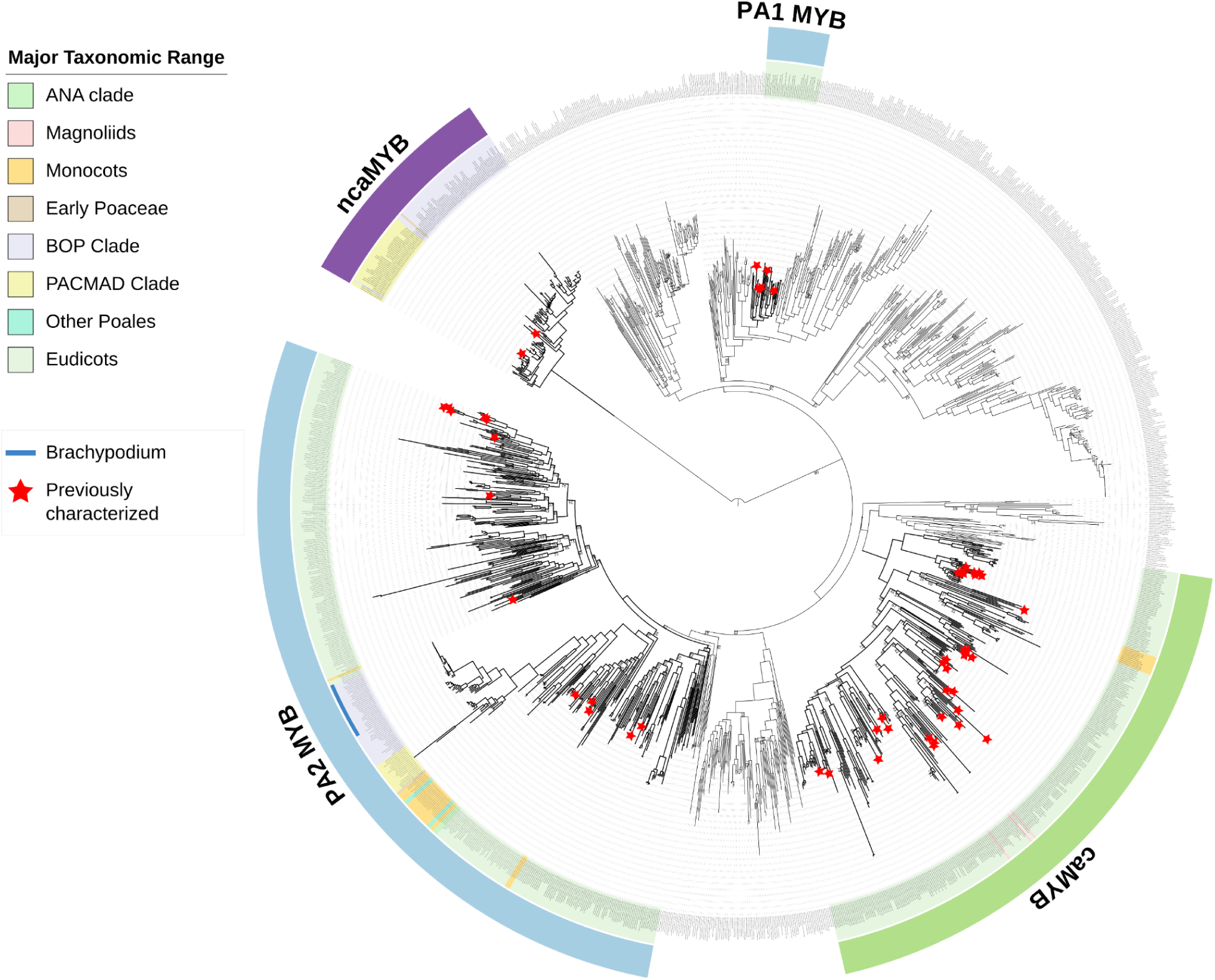
Codon-based phylogenetic analysis of anthocyanin related MYB genes. While most plants share the canonical anthocyanin MYBs (caMYB), a new kind of non-canonical anthocyanin MYB (ncaMYB) was detected only in Poaceae. Major taxonomic ranges in angiosperms are highlighted in different colours. Functional sequences characterized in previous studies are highlighted by red asterisks at the start of terminal branches. *Brachypodium* species are indicated with a narrow blue strip next to the tree labels. Bootstrap support <80% is included.

A possible explanation for the emergence of an independent anthocyanin activating MYB lineage in Poaceae could be a loss of caMYB in ancestor of grasses, and in the absence of anthocyanin activating MYBs, the ncaMYBs took over the function as a positive regulator of anthocyanin biosynthesis. To rule out a false negative result, the same analysis was repeated for proanthocyanidin activating MYBs, PA1 and PA2 MYB, close homologs of caMYB. PA2 MYB (*AtTT2* –type) is highly conserved across angiosperms, clustering with sequences from the PACMAD and BOP clades, including *Brachypodium,* thus supporting the inferred loss of the anthocyanin MYB lineage in *Poaceae*. Since the MBW complex is known to regulate a variety of plant developmental processes (Zumajo-Cardona *et al*., 2023), it is not surprising that more generic components, bHLH and WD40, are still present in canonical forms in *Poaceae*.

### 2. AN9/TT19 and BZ2 belong to independent GST lineages

Our phylogenetic analysis revealed two independent arGST lineages in association with the anthocyanin biosynthesis (**Fig. 3**). The well-characterized arGST genes *An9* (petunia) and *TT19* (arabidopsis) are located in the Phi GST clade, while BZ2 (maize) belongs to an independent and apparently *Poaceae*-specific Tau GST clade (**Fig. 3**). These two groups are not orthologous, indicating that they evolved independently to fulfill similar roles in the anthocyanin biosynthesis pathway. This relationship has been reported before, although based on a substantially smaller number of studied plant species (Alfenito *et al*., 1998). Previously, it was assumed that the transcriptional regulation of all arGST genes would have been conserved (Alfenito *et al*., 1998), but our results regarding an independent MYB lineage contradict this hypothesis. The functional arGSTs in these two independent lineages were supported by co-expression with other genes of the anthocyanin biosynthesis. While BZ2 is a *Poaceae*-specific clade, orthologs are missing in *Brachypodium*.

**Fig. 3:**
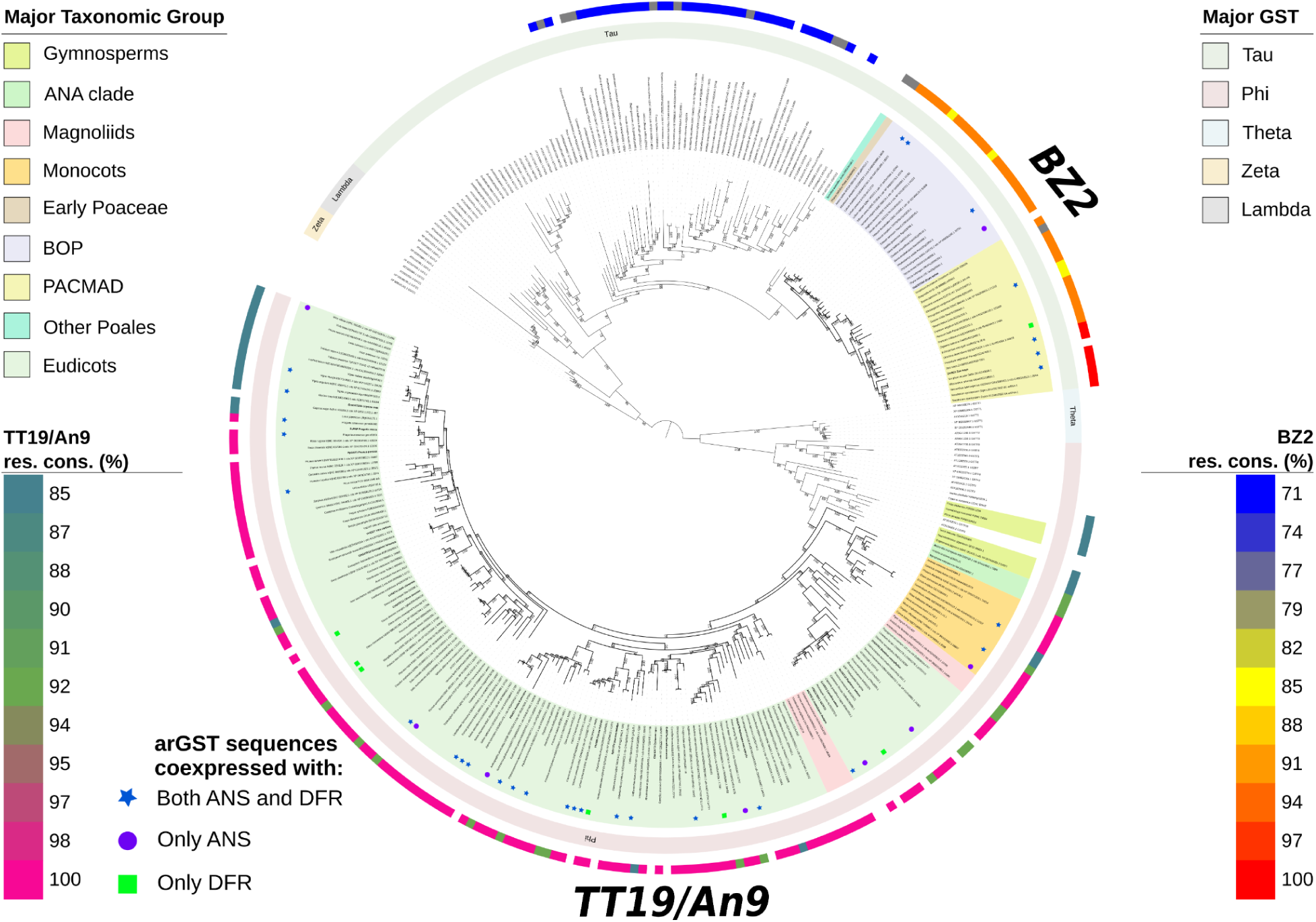
Phylogenetic relationship of landmark arGSTs. The phylogenetic tree shows two completely independent lineages of the arGSTs. The major taxonomic groups, gymnosperms, ANA, magnoliids, monocots, eudicots, BOP, and PACMAD species are denoted by green, light green, pink, yellow, light blue, light purple, and light yellow, respectively. The bold black branch labels are some of the functionally characterized arGST [LcGST4 in *Litchi chinensis* Sonn (Hu *et al*., 2016), PfGST1 in *Perilla frutescens* (Yamazaki *et al*., 2008), AN9 in *Petunia hybrida* (Lukas *et al*., 2009), DcGSTF2 in *Dianthus caryophyllus* (Sasaki *et al*., 2012), CkmGST3 in *Cyclamen* sp. (Kitamura *et al*., 2012), CsGSTF12 in *Citrus sinensis* L.(Osbeck) (Lo Piero *et al*., 2006), IbGSTF4 in *Ipomoea batatas* (Kou *et al*., 2019), GhGSTF12 in *Gossypium hirsutum* L. (Shao *et al*., 2021), FvRAP in *Fragaria vesca* (Luo *et al*., 2018), CaGSTa in *Camellia sinensis* (Liu *et al*., 2019a), PpGST1 in *Prunus persica* L. (Batsch) (Zhao *et al*., 2020), GmGST26A in *Glycine max* (Alfenito *et al*., 1998), VviGST in *Vitis vinifera* (Pérez-Díaz *et al*., 2016), HmGST9 in *Hydrangea macrophylla* (Chen *et al*., 2025), RsGSTF12 in *Raphanus sativa* (Niu *et al*., 2022), BZ2 in *Zea mays* (Marrs *et al*., 1995), OsGSTU34 in *Oryza sativa* L. (Mackon *et al*., 2024), TT19 in *Arabidopsis thaliana* (Kitamura *et al*., 2004; Sun *et al*., 2012)] and some bait sequences from KIPEs are included. The outer ring shows the major GST families, Phi, Tau, Theta, Zeta, and Lambda. The top and lower outermost rings show the percentage of conservation of functionally important amino acid residues for BZ2 and TT19/An9 function, respectively, ranging from 71% to 100 %. The blue star, purple circle and green square represents arGST sequences co-expressed with ANS and DFR, only with ANS, and only DFR, respectively.

The independent anthocyanin biosynthesis in *Poaceae* could have conferred a yet unidentified evolutionary advantage over the canonical anthocyanin biosynthesis outside the *Poaceae*. However, it is also possible that the independent recruitment of an anthocyanin regulating MYB and arGST followed a preceding loss of the anthocyanin biosynthesis. Such an anthocyanin loss could be explained by a transition phase with reduced selection pressure on a functional anthocyanin biosynthesis, e.g., due to wind pollination, as pollinator attraction is a major anthocyanin function (Davies *et al*., 2012; Grünig *et al*., 2024). A similar scenario has been proposed and discussed for the pigmentation transition in the Caryophyllales (Ehrendorfer, 1976; Brockington *et al*., 2011). Numerous anthocyanin functions besides pollinator attraction (Grünig *et al*., 2024) might explain why plant lineages without anthocyanins are rarely successful in the long term, as can be inferred from the almost universal presence of anthocyanins across recent plant lineages.

A duplication event occurred at the base of the GSTF (GST phi) in the *Brassicaceae* family, giving rise to GSTF11 and GSTF12 paralogs with two distinct functions (**Fig. S15 in Additional File 1**). The GSTF12 is an *Arabidopsis thaliana* anthocyanin-related glutathione S-transferase involved in the anthocyanin biosynthesis pathway, while GSTF11 has been postulated to be involved in the aliphatic glucosinolate biosynthesis pathway (Mikhaylova *et al*., 2021; Zhang *et al*., 2022). The *Arabidopsis thaliana* GSTF11 and GSTU20 double mutant shows a reduction in the glucosinolate contents in a dosage-dependent manner (Zhang *et al*., 2022), but GSTU20 appeared to have a more critical role than GSTF11, as evidenced by the large number of differentially expressed genes compared to GSTF11. Furthermore, the GSTF11 is also reported to be associated with plant cold stress responses and resistant to fungal infection in plants, through the balance of the glutathione homeostasis (Mikhaylova *et al*., 2021).

### 3. Complex Evolution of Proanthocyanidin Biosynthesis Pathway in Poaceae

The complete loss of LAR and unconventional ANR functionality in *Zea mays* (Lu *et al*., 2023) highlights the complexity of the proanthocyanidin biosynthesis pathway in cereals. To further investigate the evolution of proanthocyanidin biosynthesis in *Poaceae*, candidate sequence identification and conserved residue analysis of LAR and other members of the Short-chain Dehydrogenase/Reductase (SDR) enzyme family were employed. Along with *Zea mays* and *Sorghum bicolor*, all of the other members of the *Andropogoneae* tribe, as well as all of the species in the *Paniceae* tribe included in this study, lacked an apparent LAR ortholog (**Fig. 4, Fig. S7-10 in Additional File 1**). Outside of these two *Panicoideae* tribes, only *Aristida adscensionis* and *Stipagrostis hirtigluma*, members of the earliest-diverging PACMAD subfamily, *Aristidoideae*, lacked an LAR ortholog. As all of the other PACMAD species analyzed in this study have a functional LAR ortholog, these losses suggest a complex evolutionary history of LAR in PACMAD grasses.

**Fig. 4:**
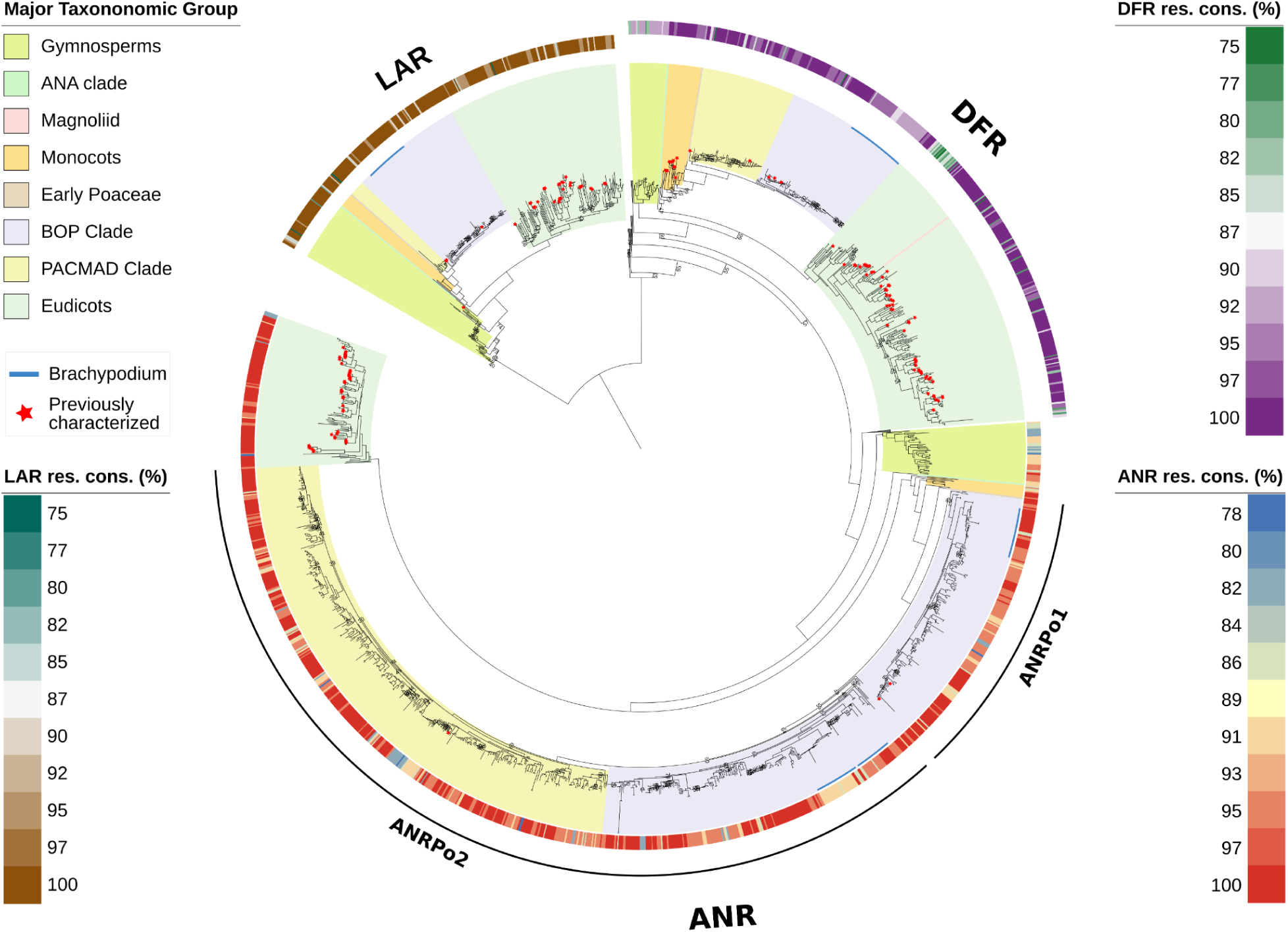
Phylogenetic relationships of Short-chain Dehydrogenase/Reductase (SDR) family enzymes involved in flavonoid biosynthesis (*DFR*, *LAR,* and *ANR*). The major taxonomic groups are highlighted in different colors. The outer ring shows the percentage of conservation of functionally important amino acid residues for all three genes in their respective functional clade. Functional sequences characterized in previous studies are highlighted by red asterisks at the start of terminal branches. Leaf labels are hidden to reduce the complexity. Bootstrap support < 80% is shown.

Another interesting evolutionary pattern revealed in this study is the formation of two distinct ANR clades in *Poaceae* following an apparent deep gene duplication event (**Fig. 4**). The first clade, which we have named ANRPo1, only contains functional gene candidates from species in the BOP clade. In contrast, the second clade, which we have named ANRPo2, contains functional gene candidates from species in both the BOP and PACMAD clades. As *Pharus latifolia*, an early-diverging Poaceae species, appears immediately outside of these clades, the gene duplication event likely occurred close to the base of the core Poaceae before the BOP and PACMAD clades diverged. The absence of paralogous ANR sequences in PACMAD species suggests lineage-specific gene loss, where, following the duplication of ANR, ANRPo1 was lost in the common ancestor of the PACMAD clade.

To gain insight into potential functional divergence between ANRPo1 and ANRPo2, expression analyses in species with both paralogs were compared using publicly available gene expression datasets (Pucker *et al*., 2025). For *Oryza nivara*, *Phyllostachys edulis*, *Lolium multiflorum*, and *Triticum urartu*, ANRPo2 shows significantly higher expression than ANRPo1. Interestingly, the opposite is true for *Brachypodium sylvaticum*, where ANRPo1 shows significantly higher expression than ANRPo2. Coexpression analysis was also performed, and both ANRPo1 and ANRPo2 were shown to be coexpressed with genes involved in abiotic stress tolerance (**Table S3, Table S4 in Additional File 1**).

### 4. Loss of anthocyanin biosynthesis in *Brachypodium sp*

Since Poaceae specific genes, ncaMYB and BZ2 were not detected in Brachypodium, we next examined the presence of all major flavonoid biosynthesis genes in the *Brachypodium* genus (*Brachypodieae* tribe). Consistent with the regulatory losses, we also failed to identify the othologs of ANS in *Brachypodium* (**Fig. S11-S14 in Additional File**). Together with the loss of ncaMYB and BZ2, the absence of ANS suggests that anthocyanin biosynthesis has been systematically lost in *Brachypodium*.

It is probable that *Poaceae*-specific anthocyanin activator ncaMYB was first lost in the *Brachypodium* genus, preventing initiation of the anthocyanin biosynthesis pathway (**Fig. 6**). Over time, the absence of anthocyanin biosynthesis activation by ncaMYB might be the driving force resulting in a subsequent loss of structural genes involved in anthocyanidin biosynthesis, namely *ANS* and *BZ2*. Interestingly, despite the loss of *ANS* in *Brachypodium* and apparent lack of cyanidins, a functional *ANR* ortholog is still retained, highlighting the complexity and unconventional nature of proanthocyanidin biosynthesis in *Poaceae*.

**Fig 5:**
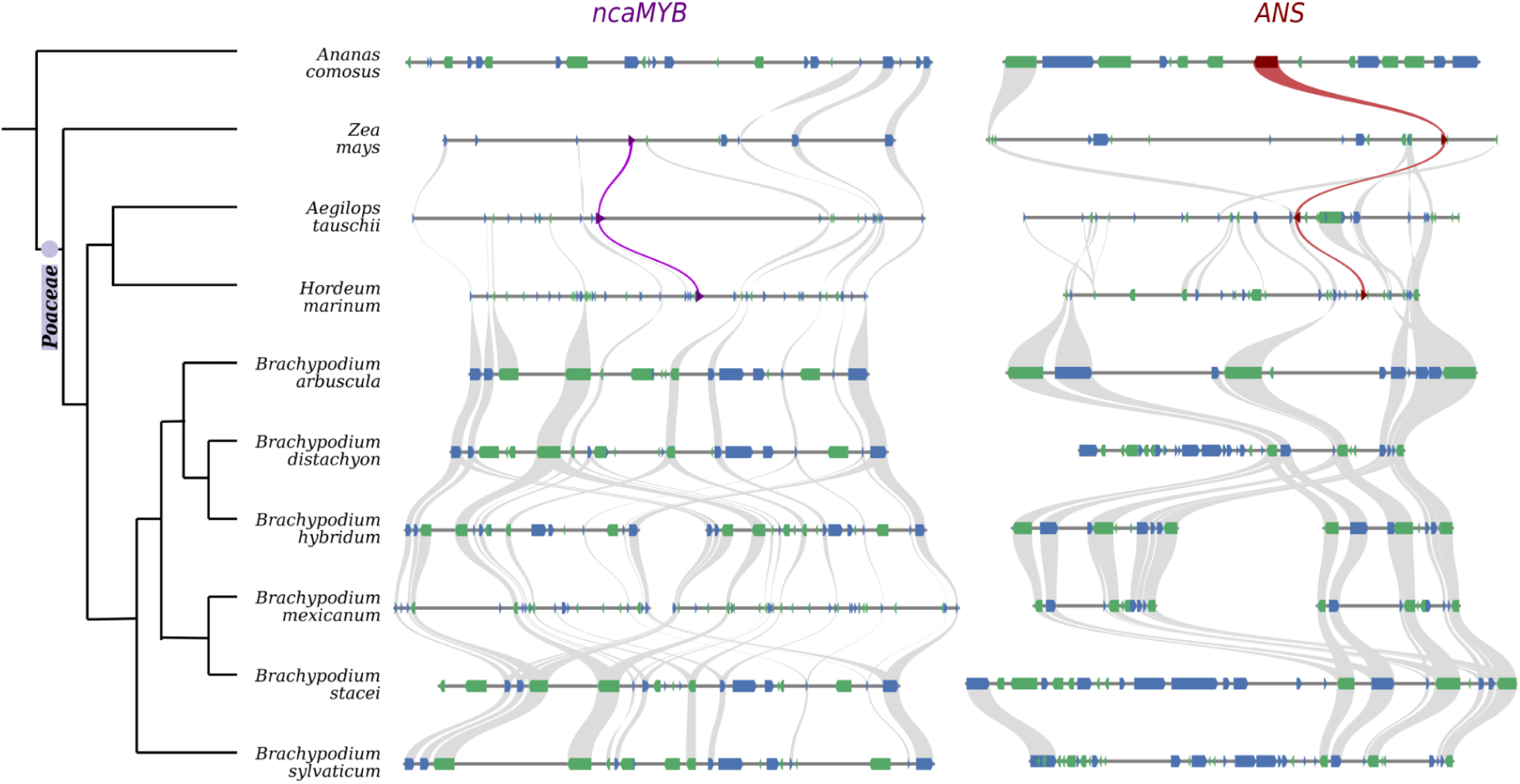
Syntenic region between *Poaceae* species and outgroup *Ananas comosus* (*Poales*) shows the specificity of the ncaMYBs to *Poaceae*, and their absence in *Brachypodium* (left). On the right, ANS microsynteny supports the loss of ANS in *Brachypodium* species.

**Fig. 6:**
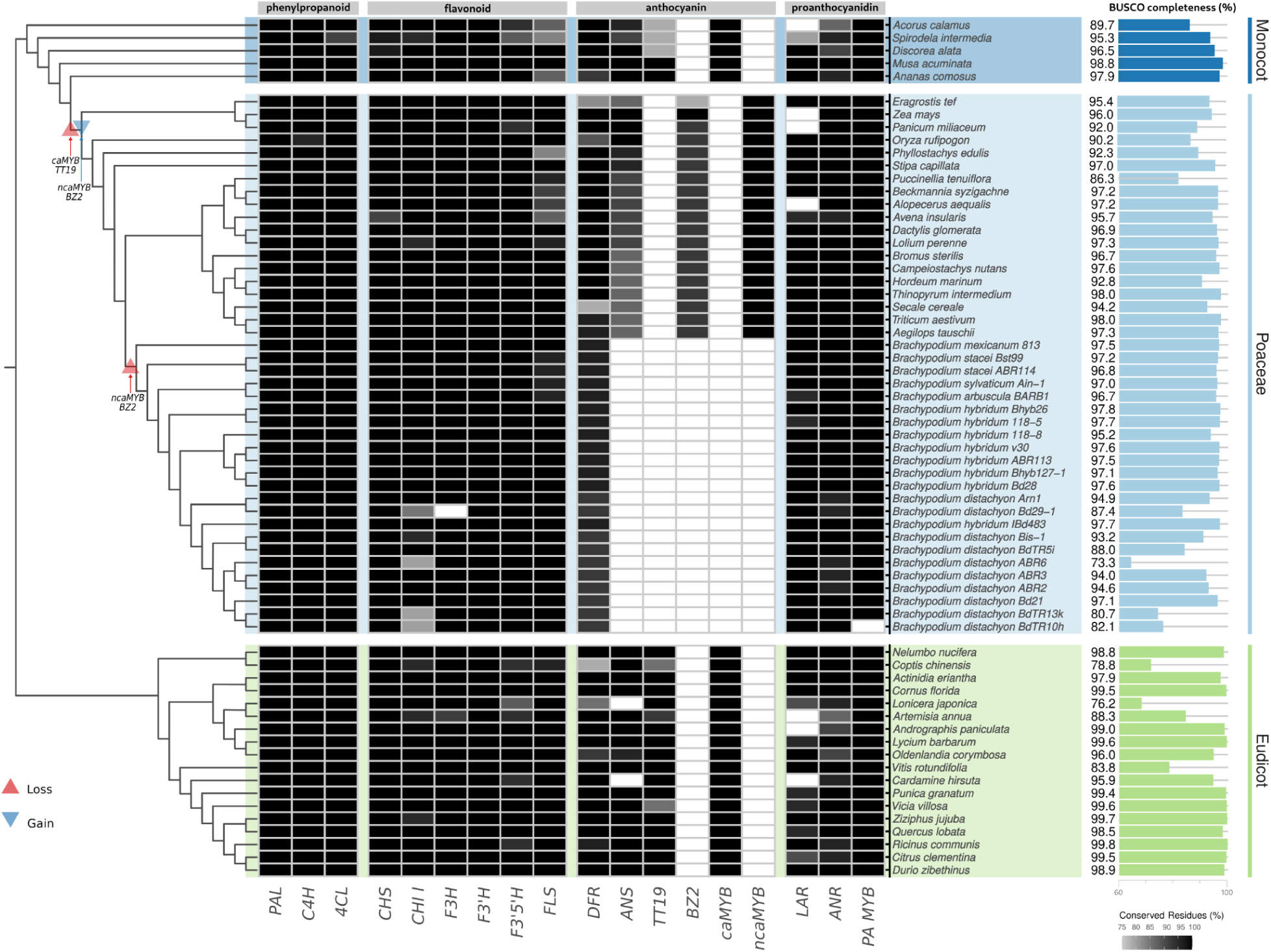
Detection of anthocyanin biosynthesis genes across angiosperms with a focus on *Poaceae*. The absence or presence of functionally conserved or putatively non-conserved genes is indicated by white, black, and grey tiles, respectively. Asterisks next to MYB123 gene label indicate the presence/absence of orthologs of *Arabidopsis thaliana* proanthocyanidin activator MYB123. Loss of anthocyanin biosynthesis genes are shown as red upwards triangles and gains as blue downwards triangles. The associated bars on the right show the BUSCO completeness (embryophyta_odb12) of the polypeptide sequence dataset used for the corresponding species in this study.

## Conclusion

The anthocyanin biosynthesis is generally considered a well-conserved pathway across land plants. Numerous anthocyanin losses reported in previous studies were restricted to the species or genus level (Marin-Recinos & Pucker, 2024). In fact, anthocyanin levels and pigmentation patterns can vary significantly within a plant species and among closely related plant species (Onozaki *et al*., 1999; Zufall & Rausher, 2003; Smith & Rausher, 2011; Marin-Recinos & Pucker, 2024; Horz *et al*., 2024). However, systematic losses of the entire anthocyanin biosynthesis in major plant lineages are rare. To the best of our knowledge, this has only been reported in some families of the core-Caryophyllales, in which anthocyanins have been replaced by betalains (Stafford, 1974, 1994; Brockington *et al*., 2011; Timoneda *et al*., 2019; Pucker *et al*., 2024b). This highlights the significance of the patterns reported here for fundamental insights into the evolution of biosynthesis pathways in plants.

## Declarations

### Data availability statement

All data sets underlying this study are publicly available (**Additional File 1**). Sequencing data have been retrieved from the Sequence Read Archive (Pucker *et al*., 2025). Customized Python scripts for the analyses in this study are available through GitHub (https://github.com/ajones292/Poaceae_ABP).

### Author contributions

NC and BP designed the project and supervised the work. NK, AJ, AR, and NC conducted analyses. AJ, AR, NC, and BP wrote the manuscript. All authors revised the manuscript and agreed to its submission.

## Acknowledgements

This work was supported by the BMBF-funded de.NBI Cloud within the German Network for Bioinformatics Infrastructure (de.NBI) (031A532B, 031A533A, 031A533B, 031A534A, 031A535A, 031A537A, 031A537B, 031A537C, 031A537D, 031A538A). We thank all members of the research group Plant Biotechnology and Bioinformatics for their discussion and support. We are particularly grateful to Milan Borchert for preliminary analyses. Open Access funding enabled and organized by Project DEAL and the University of Bonn.

