## Additional File 1 for "Evolutionary Rewiring of Anthocyanin Biosynthesis Pathway in *Poaceae*"

Khatun *et al.*

### Supplementary material

**Supplementary Table 1:** Source of genomic data sets used in this study in addition to the data submitted at 10.24355/dbbs.084-202501230512-0

| **Order** | **Family** | **Species** | **Reference** |
|---|---|---|---|
| *Acorales* | *Acoraceae* | *Acorus calamus* | (Ma, Liu, *et al.* 2023) |
| *Alismatales* | *Araceae* | *Spirodela intermedia* | (Hoang *et al.* 2020) |
| *Aquifoliales* | *Aquifoliaceae* | *Ilex paraguariensis* | (Vignale *et al.* 2025) |
| *Asparagales* | *Orchidaceae* | *Dendrobium chrysotoxum* | (Zhang, Zhang, *et al.* 2021) |
| *Asterales* | *Asteraceae* | *Erigeron canadensis* | (Laforest *et al.* 2020) |
| *Brassicales* | *Cleomaceae* | *Tarenaya hassleriana* | (Cheng *et al.* 2013) |
| *Cucurbitales* | *Datiscaceae* | *Datisca glomerata* | (Griesmann *et al.* 2018) |
| *Cupressales* | *Cupressaceae* | *Sequoiadendron giganteum* | (Scott *et al.* 2020) |
| *Cupressales* | *Taxaceae* | *Taxus baccata* | (Wickett *et al.* 2014) |
| *Dioscoreales* | *Dioscoreaceae* | *Discorea alata* | (Bredeson *et al.* 2022) |
| *Ericales* | *Ebenaceae* | *Diospyros lotus* | (Mao *et al.* 2021) |
| *Ericales* | *Ericaceae* | *Rhododendron molle* | ("Rhododendron molle genome assembly RHMOLv1") |
| *Ericales* | *Theaceae* | *Camellia sinensis* | (Lei *et al.* 2021) |
| *Fabales* | *Fabaceae* | *Lupinus luteus* | (Martinez-Hernandez *et al.* 2024) |
| *Fabales* | *Fabaceae* | *Trifolium repens* | (Kuo *et al.* 2024) |
| *Fagales* | *Fagaceae* | *Castanea mollissima* | (Wang *et al.* 2020) |
| *Lamiales* | *Plantaginaceae* | *Digitalis purpurea* | (Horz *et al.* 2024) |
| *Lamiales* | *Orobanchaceae* | *Phtheirospermum japonicum* | ("Phtheirospermum japonicum genome assembly Pjver1") |
| *Malpighiales* | *Euphorbiaceae* | *Hevea brasiliensis* | (Cheng *et al.* 2023) |
| *Malpighiales* | *Euphorbiaceae* | *Jatropha curcas* | (Jalali *et al.* 2020) |
| *Nymphaeales* | *Nymphaeaceae* | *Victoria cruziana* | (Nowak *et al.* 2025) |
| *Piperales* | *Aristolochiaceae* | *Aristolochia fimbriata* | (Qin *et al.* 2021) |
| *Piperales* | *Piperaceae* | *Piper nigrum* | (Hu *et al.* 2019) |
| *Poales* | *Poaceae* | *Aegilops tauschii* | (Luo *et al.* 2017) |
| *Poales* | *Poaceae* | *Alopecerus aequalis* | (Wright *et al.* 2024) |
| *Poales* | *Bromeliaceae* | *Ananas comosus* | (Ming *et al.* 2015) |
| *Poales* | *Poaceae* | *Avena insularis* | (Kamal *et al.* 2022) |
| *Poales* | *Poaceae* | *Beckmannia syzigachne* | (Han *et al.* 2025) |
| *Poales* | *Poaceae* | *Brachypodium arbuscula* BARB1 | ("B.arbusculaBARB1 v3.1: Phytozome") |
| *Poales* | *Poaceae* | *Brachypodium distachyon* Bd21 | (Vogel *et al.* 2010) |
| *Poales* | *Poaceae* | *Brachypodium distachyon* ABR2 | (Gordon *et al.* 2017) |

| | | | |
|---|---|---|---|
| *Poales* | *Poaceae* | *Brachypodium distachyon* ABR3 | (Gordon *et al.* 2017) |
| *Poales* | *Poaceae* | *Brachypodium distachyon* ABR6 | (Bettgenhaeuser *et al.* 2017) |
| *Poales* | *Poaceae* | *Brachypodium distachyon* Arn1 | (Gordon *et al.* 2017) |
| *Poales* | *Poaceae* | *Brachypodium distachyon* Bd29-1 | (Gordon *et al.* 2017) |
| *Poales* | *Poaceae* | *Brachypodium distachyon* BdTR10h | (Gilbert *et al.* 2018) |
| *Poales* | *Poaceae* | *Brachypodium distachyon* BdTR13k | (Gilbert *et al.* 2018) |
| *Poales* | *Poaceae* | *Brachypodium distachyon* BdTR5i | (Gordon *et al.* 2017) |
| *Poales* | *Poaceae* | *Brachypodium distachyon* Bis-1 | (Gordon *et al.* 2017) |
| *Poales* | *Poaceae* | *Brachypodium hybridum* 118-5 | (Gordon *et al.* 2020) |
| *Poales* | *Poaceae* | *Brachypodium hybridum* 118-8 | (Gordon *et al.* 2020) |
| *Poales* | *Poaceae* | *Brachypodium hybridum* ABR113 | (Gordon *et al.* 2020) |
| *Poales* | *Poaceae* | *Brachypodium hybridum* Bd28 | (Gordon *et al.* 2020) |
| *Poales* | *Poaceae* | *Brachypodium hybridum* Bhyb127-1 | (Gordon *et al.* 2020) |
| *Poales* | *Poaceae* | *Brachypodium hybridum* Bhyb26 | (Gordon *et al.* 2020) |
| *Poales* | *Poaceae* | *Brachypodium hybridum* IBd483 | (Chen *et al.* 2024) |
| *Poales* | *Poaceae* | *Brachypodium hybridum* v30 | (Gordon *et al.* 2020) |
| *Poales* | *Poaceae* | *Brachypodium mexicanum* 813 | ("B.mexicanum v2.1: BMAP") |
| *Poales* | *Poaceae* | *Brachypodium stacei* ABR114 | (Gordon *et al.* 2020) |
| *Poales* | *Poaceae* | *Brachypodium stacei* Bst99 | (Chen *et al.* 2024) |
| *Poales* | *Poaceae* | *Brachypodium sylvaticum* Ain-1 | (Lei *et al.* 2024) |
| *Poales* | *Poaceae* | *Bromus sterilis* | (Christenhusz 2024a) |
| *Poales* | *Poaceae* | *Bromus tectorum* | (Revolinski *et al.* 2023) |
| *Poales* | *Poaceae* | *Campeiostachys nutans* | (Xiong *et al.* 2025) |
| *Poales* | *Poaceae* | *Dactylis glomerata* | (Huang *et al.* 2020) |
| *Poales* | *Poaceae* | *Hordeum marinum* | (Kuang *et al.* 2022) |
| *Poales* | *Poaceae* | *Lolium perenne* | (Frei *et al.* 2021) |
| *Poales* | *Poaceae* | *Puccinellia tenuiflora* | (Guo *et al.* 2020) |
| *Poales* | *Poaceae* | *Secale cereale* | (Rabanus-Wallace *et al.* 2021) |
| *Poales* | *Poaceae* | *Stipa capillata* | (Baiakhmetov *et al.* 2021) |
| *Poales* | *Poaceae* | *Thinopyrum intermedium* | ("T.intermedium v3.1: Phytozome") |
| *Poales* | *Poaceae* | *Triticum aestivum* | (Zhu *et al.* 2021) |
| *Poales* | *Poaceae* | *Zea mays* | ("Zea mays genome assembly Zm-B73-REFERENCE-NAM-5.0") |
| *Poales* | *Poaceae* | *Aristida adscensionis* | (Pereira *et al.* 2025) |
| *Poales* | *Poaceae* | *Cenchrus macrourus* | (Zheng *et al.* 2023) |
| *Poales* | *Poaceae* | *Cenchrus purpureus* | (Yan *et al.* 2021) |
| *Poales* | *Poaceae* | *Chasmanthium latifolium* | ("Genome Warehouse") |

| | | | |
|---|---|---|---|
| *Poales* | *Poaceae* | *Coix lacryma-jobi* | (Liu *et al.* 2020) |
| *Poales* | *Poaceae* | *Dichanthelium oligosanthes* | (Studer *et al.* 2016) |
| *Poales* | *Poaceae* | *Digitaria exilis* | (Abrouk *et al.* 2020) |
| *Poales* | *Poaceae* | *Echinochloa colona* | (Wu *et al.* 2022) |
| *Poales* | *Poaceae* | *Echinochloa oryzicola* | (Wu *et al.* 2022) |
| *Poales* | *Poaceae* | *Miscanthus floridulus* | (Zhang, Ge, *et al.* 2021) |
| *Poales* | *Poaceae* | *Panicum hallii* | (Lovell *et al.* 2018) |
| *Poales* | *Poaceae* | *Panicum virgatum* | (Lovell *et al.* 2021) |
| *Poales* | *Poaceae* | *Paspalum notatum* | (Vega *et al.* 2024) |
| *Poales* | *Poaceae* | *Paspalum vaginatum* | (Sun *et al.* 2022) |
| *Poales* | *Poaceae* | *Pharus latifolius* | ("P.latifolius v1.1: Phytozome") |
| *Poales* | *Poaceae* | *Phragmites australis* | (Christenhusz 2024b) |
| *Poales* | *Poaceae* | *Saccharum longisetosum* | ("Genome Warehouse") |
| *Poales* | *Poaceae* | *Saccharum officinarum* | ("Genome Warehouse") |
| *Poales* | *Poaceae* | *Saccharum spontaneum* | (Zhang *et al.* 2018) |
| *Poales* | *Poaceae* | *Setaria italica* | (Bennetzen *et al.* 2012) |
| *Poales* | *Poaceae* | *Setaria virdis* | (Mamidi *et al.* 2020) |
| *Poales* | *Poaceae* | *Sorghum bicolor* | (Paterson *et al.* 2009) |
| *Poales* | *Poaceae* | *Sorghum virgatum* | ("Genome Warehouse") |
| *Poales* | *Poaceae* | *Stipagrostis hirtigluma* | (Pereira *et al.* 2025) |
| *Poales* | *Poaceae* | *Thysanolaena latifolia* | ("Genome Warehouse") |
| *Poales* | *Poaceae* | *Urochloa decumbens* | (Ryan *et al.* 2025) |
| *Poales* | *Poaceae* | *Avena sativa* | (Kamal *et al.* 2022) |
| *Poales* | *Poaceae* | *Pharus latifolius* | (Ma *et al.* 2021) |
| *Poales* | *Poaceae* | *Raddia guianensis* | (Guo *et al.* 2019) |
| *Poales* | *Poaceae* | *Olyra latifolia* | (Guo *et al.* 2019) |
| *Poales* | *Poaceae* | *Bonia amplexicaulis* | (Guo *et al.* 2019) |
| *Poales* | *Poaceae* | *Cleistogenes songorica* | (Zhang, Wu, *et al.* 2021) |
| *Poales* | *Poaceae* | *Eleusine coracana* | (Devos *et al.* 2023) |
| *Poales* | *Poaceae* | *Phragmites australis* | (Christenhusz *et al.* 2024) |
| *Poales* | *Poaceae* | *Digitaria radiosca* | (Minoji & Sakai 2024) |
| *Poales* | *Poaceae* | *Guadua angustifolia* | (Guo *et al.* 2019) |
| *Ranunculales* | *Ranunculaceae* | *Aquilegia coerulea* | (Filiault *et al.* 2018) |
| *Ranunculales* | *Papaveraceae* | *Papaver nudicaule* | (Catania *et al.* 2022) |
| *Sapindales* | *Sapindaceae* | *Xanthoceras sorbifolium* | (Liang *et al.* 2019) |
| *Sapindales* | *Sapindaceae* | *Dimocarpus longan* | (Lin *et al.* 2017) |

| | | | |
|---|---|---|---|
| *Sapindales* | *Nitrariaceae* | *Nitraria sibirica* | (Ma, Ru, *et al.* 2023) |
| *Saxifragales* | *Altingiaceae* | *Liquidambar formosana* | (Xu *et al.* 2024) |
| *Solanales* | *Convolvulaceae* | *Ipomoea triloba* | (Wu *et al.* 2018) |
| *Solanales* | *Solanaceae* | *Petunia axillaris* | (Bombarely *et al.* 2016) |
| *Solanales* | *Solanaceae* | *Capsicum annuum* | (Hulse-Kemp *et al.* 2018) |
| *Zingiberales* | *Musaceae* | *Musa acuminata* | (Huang *et al.* 2023) |

**Supplementary Table 2:** Gene prediction statistics of the 13 genomes annotated in this study.

| Species | No. of genes anotated | No. of transcripts annotated | Protein BUSCO (embryophyta_odb12) |
|---|---|---|---|
| *Stipa capillata* | 59070 | 78347 | 99.20% |
| *Brachypodium hybridum* 118 | 62461 | 81736 | 99.40% |
| *Brachypodium distachyon* ABR6 | 29866 | 35598 | 73.30% |
| *Brachypodium distachyon* BdTR13k | 30754 | 36788 | 80.70% |
| *Brachypodium distachyon* BdTR10h | 31020 | 37230 | 82.10% |
| *Brachypodium hybridum* Bhyb127-*1* | 65611 | 74134 | 97.70% |
| *Brachypodium hybridum* IBd483 | 65533 | 74687 | 97.70% |
| *Brachypodium stacei* Bst99 | 32749 | 39581 | 97.20% |
| *Bromus sterilis* | 46379 | 60914 | 98.70% |
| *Puccinellia tenuiflora* | 48326 | 50477 | 86.30% |
| *Bromus tectorum* | 43477 | 58028 | 91.70% |
| *Beckmannia syzigachne* | 59299 | 74372 | 97.20% |
| *Dactylis glomerata* | 48254 | 63151 | 98.90% |

**Supplementary Table 3:** Top coexpressed gene candidates with sequences from ANRPo1 clade. The model plant *Arabidopsis thaliana* was used to tranfer annotation based on best match candidates.

| **Gene ID** | **Symbol** | **Description** |
|---|---|---|
| AT1G01770 | propionyl-CoA carboxylase | Gene |
| AT1G06330 | Heavy metal transport/detoxification superfamily protein | Gene |
| AT1G14900 | HMGA | high mobility group A |
| AT1G29280 | WRKY65 | WRKY DNA-binding protein 65 |
| AT1G59960 | NAD(P)-linked oxidoreductase superfamily protein | Gene |
| AT2G03200 | Eukaryotic aspartyl protease family protein | Gene |
| AT2G16290 | F-box SKIP23-like protein (DUF295) | Gene |
| AT2G16770 | bZIP23 | Basic-leucine zipper (bZIP) transcription factor family |
| AT2G32300 | UCC1 | uclacyanin 1 |
| AT3G04890 | adenine phosphoribosyltransferase-like protein, putative (DUF2358) | Gene |
| AT3G07040 | RPM1 | NB-ARC domain-containing disease resistance protein |
| AT3G14460 | LRR and NB-ARC domains-containing disease resistance protein | Gene |
| AT3G14470 | NB-ARC domain-containing disease resistance protein | Gene |
| AT3G18830 | PMT5 | polyol/monosaccharide transporter 5 |
| AT3G47570 | Leucine-rich repeat protein kinase family protein | Gene |
| AT3G48140 | B12D protein | Gene |
| AT4G05200 | CRK25 | cysteine-rich RLK (RECEPTOR-like protein kinase) 25 |
| AT4G23180 | CRK10 | cysteine-rich RLK (RECEPTOR-like protein kinase) 10 |
| AT4G30110 | HMA2 | heavy metal atpase 2 |
| AT5G23850 | O-glucosyltransferase rumi-like protein (DUF821) | Gene |

**Supplementary Table 4:** Top coexpressed gene candidates with sequences from *ANRPo2* clade. The model plant *Arabidopsis thaliana* was used to tranfer annotation based on best match candidates.

| **Gene ID** | **Symbol** | **Description** |
|---|---|---|
| AT1G02335 | GL22 | germin-like protein subfamily 2 member 2 precursor |
| AT1G05260 | RCI3 | Peroxidase superfamily protein |
| AT1G06840 | Leucine-rich repeat protein kinase family protein | Gene |
| AT1G10360 | GSTU18 | glutathione S-transferase TAU 18 |
| AT1G10370 | ERD9 | Glutathione S-transferase family protein |
| AT1G15690 | AVP1 | Inorganic H pyrophosphatase family protein |
| AT1G15950 | CCR1 | cinnamoyl coa reductase 1 |
| AT1G30900 | VSR6 | VACUOLAR SORTING RECEPTOR 6 |
| AT1G47290 | 3BETAHSD/D1 | 3beta-hydroxysteroid-dehydrogenase/decarboxylase isoform 1 |
| AT1G61250 | SC3 | secretory carrier 3 |
| AT1G61720 | BAN | NAD(P)-binding Rossmann-fold superfamily protein |
| AT1G62300 | WRKY6 | WRKY family transcription factor |
| AT1G65680 | EXPB2 | expansin B2 |
| AT1G65870 | Disease resistance-responsive (dirigent-like protein) family protein | Gene |
| AT1G68530 | KCS6 | 3-ketoacyl-CoA synthase 6 |
| AT2G18360 | alpha/beta-Hydrolases superfamily protein | Gene |
| AT2G18750 | Calmodulin-binding protein | Gene |
| AT2G35270 | GIK | Putative AT-hook DNA-binding family protein |
| AT2G36530 | LOS2 | Enolase |
| AT2G37710 | RLK | receptor lectin kinase |
| AT2G38760 | ANNAT3 | annexin 3 |
| AT2G39420 | alpha/beta-Hydrolases superfamily protein | Gene |
| AT2G45910 | U-box domain-containing protein kinase family protein | Gene |
| AT2G46950 | CYP709B2 | cytochrome P450, family 709, subfamily B, polypeptide 2 |
| AT2G47890 | B-box type zinc finger protein with CCT domain-containing protein | Gene |
| AT3G03800 | SYP131 | syntaxin of plants 131 |
| AT3G07040 | RPM1 | NB-ARC domain-containing disease resistance protein |
| AT3G09270 | GSTU8 | glutathione S-transferase TAU 8 |

| | |
|---|---|
| AT3G14310 PME3 | pectin methylesterase 3 |
| AT3G14360 alpha/beta-Hydrolases superfamily protein | Gene |
| AT3G14460 LRR and NB-ARC domains-containing disease resistance protein | Gene |
| AT3G14470 NB-ARC domain-containing disease resistance protein | Gene |
| AT3G16520 UGT88A1 | UDP-glucosyl transferase 88A1 |
| AT3G47570 Leucine-rich repeat protein kinase family protein | Gene |
| AT3G48310 CYP71A22 | cytochrome P450, family 71, subfamily A, polypeptide 22 |
| AT3G53150 UGT73D1 | UDP-glucosyl transferase 73D1 |
| AT3G53810 Concanavalin A-like lectin protein kinase family protein | Gene |
| AT3G59990 MAP2B | methionine aminopeptidase 2B |
| AT4G01950 GPAT3 | glycerol-3-phosphate acyltransferase 3 |
| AT4G05200 CRK25 | cysteine-rich RLK (RECEPTOR-like protein kinase) 25 |
| AT4G09320 NDPK1 | nucleoside diphosphate kinase |
| AT4G18360 GOX3 | Aldolase-type TIM barrel family protein |
| AT4G20820 FAD-binding Berberine family protein | Gene |
| AT4G23180 CRK10 | cysteine-rich RLK (RECEPTOR-like protein kinase) 10 |
| AT4G38690 PLC-like phosphodiesterases superfamily protein | Gene |
| AT4G39490 CYP96A10 | cytochrome P450, family 96, subfamily A, polypeptide 10 |
| AT4G40070 RING/U-box superfamily protein | Gene |
| AT5G05860 UGT76C2 | UDP-glucosyl transferase 76C2 |
| AT5G06740 Concanavalin A-like lectin protein kinase family protein | Gene |
| AT5G06839 TGA10 | bZIP transcription factor family protein |
| AT5G13080 WRKY75 | WRKY DNA-binding protein 75 |
| AT5G39670 Calcium-binding EF-hand family protein | Gene |
| AT5G39785 hypothetical protein (DUF1666) | Gene |
| AT5G41410 BEL1 | POX (plant homeobox) family protein |
| AT5G45800 MEE62 | Leucine-rich repeat protein kinase family protein |
| AT5G57800 CER3 | Fatty acid hydroxylase superfamily |
| AT5G66550 Maf-like protein | Gene |

**Supplementary Table 5:** RNA-seq IDs used for co-expression analysis

| | |
|---|---|
| *Phragmites australis* | SRR16647505,SRR16647506,SRR16647507,SRR16647508,SRR16647509,SRR16647510,SRR16647511,SRR16647512,SRR16647513,SRR16647514,SRR16647515,SRR16647516,SRR16647517,SRR16647518,SRR16647519,SRR16647520,SRR16647521,SRR16647522,SRR16647523,SRR16647524,SRR16647525,SRR16647526,SRR16647527,SRR16647528,SRR16647529,SRR16647530,SRR16647531,SRR16647532,SRR16647533,SRR16647534,SRR16647535,SRR16647536,SRR16647537,SRR16647538,SRR16647539,SRR16647540,SRR16647541,SRR16647542,SRR16647543,SRR16647544,SRR16647545,SRR16647546,SRR16647547,SRR16647548,SRR16647549,SRR16647550,SRR16647551,SRR16647552,SRR16647553,SRR16647554,SRR16647555,SRR16647556,SRR16647557,SRR16647558,SRR16647559,SRR16647560,SRR16647561,SRR16647562,SRR16647563,SRR16647564,SRR16647565,SRR16647566,SRR16647567,SRR16647568,SRR16647569,SRR16647570,SRR16647571,SRR16647572,SRR16647573,SRR16647574,SRR16647575,SRR16647576,SRR16647577,SRR16647578,SRR16647579,SRR16647580,SRR16647581,SRR16647582,SRR16647583,SRR16647584,SRR16647585,SRR16647586,SRR16647587,SRR16647588,SRR16647589,SRR16647590,SRR16647591,SRR16647592,SRR16647593,SRR16647594,SRR16647595,SRR16647596,SRR16647597,SRR16647598,SRR16647599,SRR16647600,SRR16647601,SRR16647602,SRR16647603,SRR16647604,SRR16647605,SRR16647606,SRR16647607,SRR16647608,SRR16647609,SRR16647610,SRR16647611,SRR16647612,SRR16647613,SRR16647614,SRR16647615,SRR16647616,SRR16647617,SRR16647618,SRR16647619,SRR16647620,SRR16647621,SRR16647622,SRR16647623,SRR16647624,SRR16647625,SRR16647626,SRR16647627,SRR16647628,SRR16647629,SRR16647630,SRR16647631,SRR16647632,SRR16647633,SRR16647634,SRR16647635,SRR16647636,SRR16647637,SRR16647638,SRR16647639,SRR16647640,SRR16647641,SRR16647642,SRR16647643,SRR16647644,SRR16647645,SRR16647646,SRR16647647,SRR16647648,SRR16647649,SRR16647650,SRR16647651,SRR16647652,SRR16647653,SRR16647654,SRR16647655,SRR16647656,SRR16647657,SRR16647658,SRR16647659,SRR16647660,SRR16647661,SRR16647662,SRR16647663,SRR16647664,SRR16647665,SRR16647666,SRR16647667,SRR16647668,SRR16647669,SRR16647670,SRR16647671,SRR16647672,SRR16647673,SRR16647674,SRR16647675,SRR16647676,SRR16647677,SRR16647678,SRR16647679,SRR16647680,SRR16647681,SRR16647682,SRR16647683,SRR16647684,SRR16647685,SRR16647686,SRR16647687,SRR16647688,SRR16647689,SRR16647690,SRR16647691,SRR16647692,SRR16647693,SRR16647694,SRR16647695,SRR16647696,SRR16648326,SRR16648327,SRR16648328,SRR16648329,SRR16648330,SRR16648331,SRR16648332,SRR19646716,SRR19646717,SRR19646718,SRR19646719,SRR19646720,SRR19646721,SRR19646722,SRR3212855,SRR3233385,SRR3233386,SRR3233387,SRR3233388,SRR3233389,SRR3233390,SRR3233391,SRR3233392,SRR3233393,SRR3233394,SRR3233395,SRR3233396,SRR3233397,SRR3233398,SRR15852918,SRR15852919,SRR15852920,SRR15852921,SRR13300084,SRR13300085,SRR13300086,SRR13300087,SRR13300088,SRR13300089,SRR15783382,SRR9648369,SRR9648370,SR |

| | |
|---|---|
| | R9648371,SRR9648372,SRR9648373,SRR9648374,SRR9648375,SRR9648376,SRR9648377,SRR9648378,SRR9648379,SRR9648380,SRR26557086,SRR26557087,SRR26557088,SRR26557089,SRR26557090,SRR26557091,SRR15902994,ERR3891548 |
| *Eleusine coracana* | SRR1663469,SRR1663470,SRR3977085,SRR3146431,SRR3146438,SRR5341138,SRR5341139,SRR5341140,SRR5341141,SRR5341142,SRR5341143,SRR5341144,SRR5341145,SRR5341146,SRR5341147,SRR5341148,SRR6984602,SRR6984636,SRR1038430,SRR1038431,SRR1038432,SRR1038433,SRR1038434,SRR1038435,SRR1038436,SRR1038437,SRR2001609,SRR2001830,SRR4021829,SRR4021830,SRR19328872,SRR19328873,SRR19328874,SRR19328875,SRR19328876,SRR19328877,SRR12320327,SRR12320328,SRR12320329,SRR12320330,SRR12320331,SRR12320332,SRR12320333,SRR12320334,SRR12320335,SRR12320336,SRR12320337,SRR12320338,SRR12320339,SRR12320340,SRR12320341,SRR12320342,SRR12320343,SRR12320344,SRR12320345,SRR12320346,SRR12320347,SRR12320348,SRR12320349,SRR12320350,SRR12320351,SRR12320352,SRR12320353,SRR12320354,SRR10948829,SRR10948830,SRR10948831,SRR10948832,SRR3156038,SRR3156044,ERR2040786,ERR2040787,ERR2040788,ERR2040789,SRR7223711,SRR7223712,SRR5468572,SRR5468573,SRR1151079,SRR1151080 |
| *Nitraria sibirica* | SRR23703247,SRR23703248,SRR23703249,SRR23703250,SRR23703251,SRR23703252,SRR23703253,SRR23703254,SRR23703255,SRR22403917,SRR22403918,SRR22403919,SRR22403920,SRR22403921,SRR22403922,SRR23683940,SRR23683942,SRR23683943,SRR23683944,SRR23683945,SRR23683946,SRR23683947,SRR23683950,SRR23683951,SRR23683952,SRR23683953,SRR23683954,SRR23683955,SRR23683956,SRR23683958,SRR23683959,SRR23683962,SRR23683965,SRR23683966,SRR23683970,SRR23683971,SRR23683941,SRR23683948,SRR23683949,SRR23683957,SRR23683960,SRR23683963,SRR23683964,SRR23683967,SRR23683968,SRR23683969,SRR13417509,SRR13417510,SRR13417511,SRR13417512,SRR13417513,SRR13417514,SRR13417515,SRR13417516,SRR13417517,SRR13417518,SRR13417519,SRR13417520,SRR7013767,SRR7013768,SRR7013769,SRR7013770,SRR7013771,SRR7013772,SRR7013773,SRR7013774,SRR7013775,SRR5585869,SRR10829649,SRR10829663,SRR10829664,SRR10829657,SRR10829658,SRR10829659,SRR10829660,SRR10829661,SRR10829662 |
| *Dimocarpus longan* | SRR23892005,SRR23892006,SRR23892004,SRR23825259,SRR23825260,SRR23825261,SRR412534,SRR3925166,SRR3925167,SRR3925168,SRR3925169,SRR3925170,SRR3925171,SRR3925172,SRR3925173,SRR3925174,SRR3925175,SRR3925176,SRR3925177,SRR3925178,SRR3925179,SRR27104555,SRR27104556,SRR27104557,SRR27104558,SRR27104559,SRR27104560,SRR27104561,SRR27104562,SRR27104563,SRR7614757,SRR7614758,SRR7614759,SRR7614760,SRR7614761,SRR7614762,SRR9673357,SRR9673358,SRR9673359,SRR9673360,SRR9673361,SRR9673362,SRR9673363,SRR9673364,SRR9673365,SRR9673366,SRR9673367,SRR9673368, |

| | |
|---|---|
| | SRR9673369,SRR9673370,SRR9673371,SRR9673372,SRR9673373,SRR9673374,SRR14978271,SRR14978272,SRR14978273,SRR14978274,SRR14978275,SRR14978276,SRR14978277,SRR14978278,SRR14978279,SRR14978280,SRR14978281,SRR14978282,SRR14978283,SRR14978284,SRR14978285,SRR14978286,SRR14978287,SRR14978288,SRR14978289,SRR14978290,SRR14978291,SRR14978292,SRR14978293,SRR14978294,SRR14978295,SRR14978296,SRR14978297,SRR14978298,SRR14978299,SRR14978300,SRR14978301,SRR14978302,SRR14978303,SRR14978304,SRR14978305,SRR14978306,SRR14978307,SRR14978308,SRR14978309,SRR14978310,SRR14978311,SRR14978312,SRR12641085,SRR12641086,SRR12042863,SRR12042864,SRR12042865,SRR12042866,SRR12042867,SRR12042868,SRR12042869,SRR12042870,SRR12042871,SRR12042872,SRR12042873,SRR12042874,SRR12042875,SRR12042876,SRR12042877,SRR12042878,SRR12042879,SRR12042880,SRR12042881,SRR12042882,SRR12042883,SRR12042884,SRR12042885,SRR12042886,SRR12042887,SRR12042888,SRR12042889,SRR12042890,SRR12042891,SRR12042892,SRR10112229,SRR10112230,SRR10112231,SRR10112232,SRR15364625,SRR15364626,SRR15364627,SRR15364628,SRR15364629,SRR15364630,SRR15364631,SRR15364632,SRR15364633,SRR15364634,SRR15364635,SRR15364636,SRR15364637,SRR15364638,SRR15364639,SRR15364640,SRR15364641,SRR15364642,SRR15364643,SRR15364644,SRR15364645,SRR15364646,SRR15364647,SRR15364648,SRR15364649,SRR15364650,SRR15364651,SRR15364652,SRR15364653,SRR15364654,SRR15364655,SRR15364656,SRR15364657,SRR15364658,SRR15364659,SRR15364660,SRR15364661,SRR15364662,SRR15364663,SRR15364664,SRR15364665,SRR15364666,SRR15364667,SRR15364668,SRR15364669,SRR15364670,SRR15364671,SRR15364672,SRR15364673,SRR15364674,SRR15364675,SRR15364676,SRR15364677,SRR15364678,SRR15364679,SRR15364680,SRR15364681,SRR15364682,SRR15364683,SRR15364684,SRR15367959,SRR15367961,SRR15367962,SRR15367963,SRR15367965,SRR15367966,SRR15367967,SRR15367968,SRR15367970,SRR15367971,SRR15367972,SRR19213574,SRR19213575,SRR19213576,SRR19213577,SRR19213578,SRR19213579,SRR18900563,SRR18900564,SRR18900565,SRR18900566,SRR18900567,SRR18900568,SRR18900569,SRR18900570,SRR18900571,SRR18900572,SRR18900573,SRR18900574,SRR18900575,SRR18900576,SRR18900577,SRR22064614,SRR22064615,SRR2864836,SRR5630881,SRR5630882,SRR5630883,SRR5630884,SRR5630885,SRR5630886,SRR5630887,SRR5630888,SRR5630889,SRR5630890,SRR5630891,SRR5630892,SRR5630893,SRR5630894,SRR5630895,SRR5630896,SRR5630897,SRR5630898,SRR1180011,SRR1182434,SRR3327320,SRR14654172,SRR14654173,SRR22942899,SRR22942900,SRR22942901,SRR22942902,SRR22942903,SRR22942904,SRR22942905,SRR22942906,SRR24474165,SRR24474166,SRR24474167,SRR24474168,SRR24474169,SRR24474170,SRR24474171,SRR24474172,SRR24474173 |
| *Xanthoceras sorbifolium* | SRR23148964,SRR23148965,SRR23148966,SRR23148967,SRR23148968,SRR23148969,SRR23148970,SRR23148971,SRR23148972,SRR8237443,SRR8237444,SRR8237445,SRR8237446,SRR8237447,SRR8237448,SRR8237449,SRR8237450,SRR8237451,SRR8237452,SRR8237453,SRR8237454,SRR8237455,SRR8237456,SRR8237457,SRR8237458,SRR8647435,SRR8647436,SRR8647437,SRR8647438,SRR8647439,SRR8647440,SRR8647441, |

SRR8647442,SRR8647443,SRR8647444,SRR13624816,SRR13624817,SRR13624818,SRR13624819,SRR13624820,SRR13624821,SRR13624822,SRR13624823,SRR13624824,SRR13624825,SRR13624826,SRR13624827,SRR13624828,SRR13624829,SRR13624830,SRR13624831,SRR13624832,SRR13624833,SRR13624834,SRR13624835,SRR13624836,SRR13624837,SRR13624838,SRR13624839,SRR13624840,SRR13624841,SRR13624842,SRR13624843,SRR13624844,SRR13624845,SRR13624846,SRR13624847,SRR13624848,SRR13624849,SRR13624850,SRR13624851,SRR13624852,SRR13624853,SRR13624854,SRR13624855,SRR13624856,SRR13624857,SRR13624858,SRR13624859,SRR13624860,SRR13624861,SRR13624862,SRR13624863,SRR13624864,SRR13624865,SRR13624866,SRR13624867,SRR13624868,SRR13624869,SRR13624870,SRR13624871,SRR13624872,SRR13624873,SRR13624874,SRR13624875,SRR13624876,SRR13624877,SRR13624878,SRR13624879,SRR13624880,SRR13624881,SRR13624884,SRR13624885,SRR13624886,SRR13624887,SRR13624888,SRR13624889,SRR13624890,SRR13624891,SRR13624892,SRR13624893,SRR13624894,SRR13624895,SRR13624896,SRR13624897,SRR13624898,SRR13624899,SRR13624900,SRR13624901,SRR13624902,SRR13624903,SRR13624904,SRR13624905,SRR13624906,SRR13624907,SRR13624908,SRR13624909,SRR13624910,SRR13624911,SRR13624912,SRR13624913,SRR13624914,SRR13624915,SRR13624916,SRR13624917,SRR13624918,SRR13624919,SRR13624920,SRR13624921,SRR13624922,SRR13624923,SRR13624924,SRR13624925,SRR13624926,SRR13624927,SRR13624928,SRR13624929,SRR13624930,SRR13624931,SRR13624932,SRR13624933,SRR13624934,SRR13624935,SRR13624936,SRR13624937,SRR13624938,SRR13624939,SRR13624940,SRR13624941,SRR13624942,SRR13624943,SRR13624944,SRR13624945,SRR13624946,SRR13624947,SRR13624948,SRR13624949,SRR13624950,SRR13624951,SRR13624952,SRR13624953,SRR13624954,SRR13624955,SRR13624956,SRR13624957,SRR13624958,SRR13624959,SRR13624961,SRR13624962,SRR13624963,SRR13624964,SRR13624965,SRR13624966,SRR13624967,SRR13624968,SRR13624969,SRR13624970,SRR13624972,SRR13624973,SRR13624974,SRR13624975,SRR13624976,SRR13624977,SRR13624978,SRR13624979,SRR13624980,SRR13624981,SRR13624982,SRR13624983,SRR13624984,SRR13624985,SRR13624986,SRR13624987,SRR13624988,SRR13624989,SRR13624990,SRR13624991,SRR14470652,SRR14470653,SRR14470654,SRR14470655,SRR14470656,SRR14470657,SRR14470658,SRR14470659,SRR14470660,SRR10572751,SRR10572752,SRR10572753,SRR10572754,SRR10572755,SRR10572756,SRR11217892,SRR11217893,SRR11217894,SRR11217895,SRR11217896,SRR11217897,SRR11217898,SRR11217899,SRR11217900,SRR11217901,SRR11217902,SRR11217903,SRR13038199,SRR13038200,SRR13038201,SRR13038202,SRR13038203,SRR13038204,SRR13038205,SRR13038206,SRR13038207,SRR13038208,SRR13038209,SRR13038210,SRR15696770,SRR15696771,SRR15696772,SRR15696773,SRR15696774,SRR15696775,SRR13781843,SRR13781844,SRR13781845,SRR13781846,SRR13781847,SRR13781848,SRR13781849,SRR13781850,SRR12415588,SRR12415589,SRR12415590,SRR12415591,SRR12415592,SRR12415593,SRR12415594,SRR12415595,SRR12415596,SRR12415597,SRR12415598,SRR12415599,SRR12415600,SRR12415601,SRR12415602,SRR22474645,SRR22474646,SRR22474647,SRR22474648,SRR22474649,SRR22474650,SRR22669217,SRR22669218,SRR22669219,SRR22669220,SRR22669221,SRR22669222,SRR23071

| | |
|---|---|
| | 991,SRR23071992,SRR23071993,SRR23071994,SRR23071995,SRR23071996,SRR23071997,SRR23071998,SRR23071999,SRR23072000,SRR23072001,SRR23072002,SRR23072003,SRR23072004,SRR23072005,SRR23072006,SRR23072007,SRR23072008 |
| *Lupinus luteus* | SRR19882919,SRR19882920,SRR19882925,SRR19882926,SRR19882927,SRR19882930,SRR19882931,SRR19882932,SRR19882935,SRR19882936,SRR19882937,SRR19882938,SRR19882939,SRR19882940,SRR19882918,SRR19882921,SRR19882922,SRR19882923,SRR19882924,SRR19882928,SRR19882929,SRR19882933,SRR19882934,SRR5828111,SRR5828112,SRR5828118,SRR2074729,SRR2075028,SRR2075590,SRR2075591,SRR2075594,SRR2075595,SRR2075857,SRR2075858,SRR28342699,SRR28342700,SRR28342701,SRR28342702,SRR28342703,SRR28342704,SRR28342705,SRR28342706,SRR28342707,SRR28342708,SRR28342709,SRR6317769,SRR6317770,SRR6317771,SRR6317772,SRR6317773,SRR6317774,SRR6317775,SRR6317776,SRR6317786,SRR6317787,SRR6317788,SRR6317789,SRR6317790,SRR6317791,SRR6317792,SRR6317793,SRR6317794,SRR6317795,SRR1745794,SRR5828115,SRR5828116,SRR5828117,SRR2076751,SRR2076752,SRR2078878,SRR2078879,SRR21902491,SRR21902492,SRR30893544,SRR30893545,SRR30893546,SRR30893547,SRR30893548,SRR30893549,SRR30893550,SRR30893551,SRR30893552,SRR30893553,SRR30893554,SRR30893555,SRR30893556,SRR30893557,SRR30893558,SRR30893559,SRR30893560,SRR30893561,SRR30893562,SRR30893563,SRR30893564,SRR30893565,SRR30893566,SRR30893567,SRR30893568,SRR30893569,SRR30893570,SRR30893571,SRR30893572,SRR30893573,SRR30893574,SRR30893575 |
| *Petunia axillaris* | SRR10416215,SRR10416216,SRR10416217,SRR10416218,SRR10416221,SRR10416222,SRR15174992,SRR15278148,SRR15278149,SRR15278150,SRR15278151,SRR15278152,SRR15278153,SRR2874476,SRR2874498,SRR2874980,SRR1585615,SRR1585635,SRR1585830,SRR1585954,SRR1585955,SRR8644904,SRR8644905,SRR8644906,SRR8644907,SRR8644908,SRR8644909,SRR8644910,SRR8644911,SRR8644912,SRR8644913,SRR8644914,SRR8644915,SRR17617419,SRR17617420,SRR17617421,SRR17617386,SRR17617387,SRR17617388,SRR17617389,SRR17617390,SRR17617391,SRR17617392,SRR17617393,SRR17617394,SRR17617395,SRR17617396,SRR17617397,SRR17617398,SRR17617399,SRR17617400,SRR17617401,SRR17617402,SRR17617403,SRR17617404,SRR17617405,SRR17617406,SRR17617407,SRR17617408,SRR17617409,SRR17617410,SRR17617411,SRR17617412,SRR17617413,SRR17617414,SRR17617415,SRR17617416,SRR17617417,SRR17617418,SRR21570256,SRR21570285 |
| *Piper nigrum* | ERR4099794,ERR4099795,ERR4099796,ERR4099797,ERR4099798,ERR4099799,ERR4099800,ERR4099801,ERR4099802,ERR4099803,ERR4099804,ERR4099805,SRR14876425,SRR14876426,SRR14876427,SRR14876428,SRR14876429,SRR14876430,SRR14876431,SRR14876432,SRR14876433,SRR14876434,SRR1776865,SRR1777719,SRR1781514,SRR21197322,SRR21655382,SRR21655383,SRR21655384,SRR21655385,SRR21655386,SR |

R21655387,SRR31392206,SRR31392207,SRR31392208,SRR3404020,SRR3404021,SRR3404389,SRR3404390,SRR3404569,SRR3406929,SRR408047,SRR8446741,SRR8446742,SRR8816469,SRR8816470,SRR8816471,SRR8816472,SRR8816473,SRR8816474,SRR8816475,SRR8816476,SRR8816477,SRR8816478,SRR8816480,SRR8816481,SRR8816482,SRR8816483,SRR8816484,SRR8816485,SRR8816486,SRR8816487,SRR8816488,SRR8816489,SRR8816490,SRR8816491,SRR8816492,SRR8816493

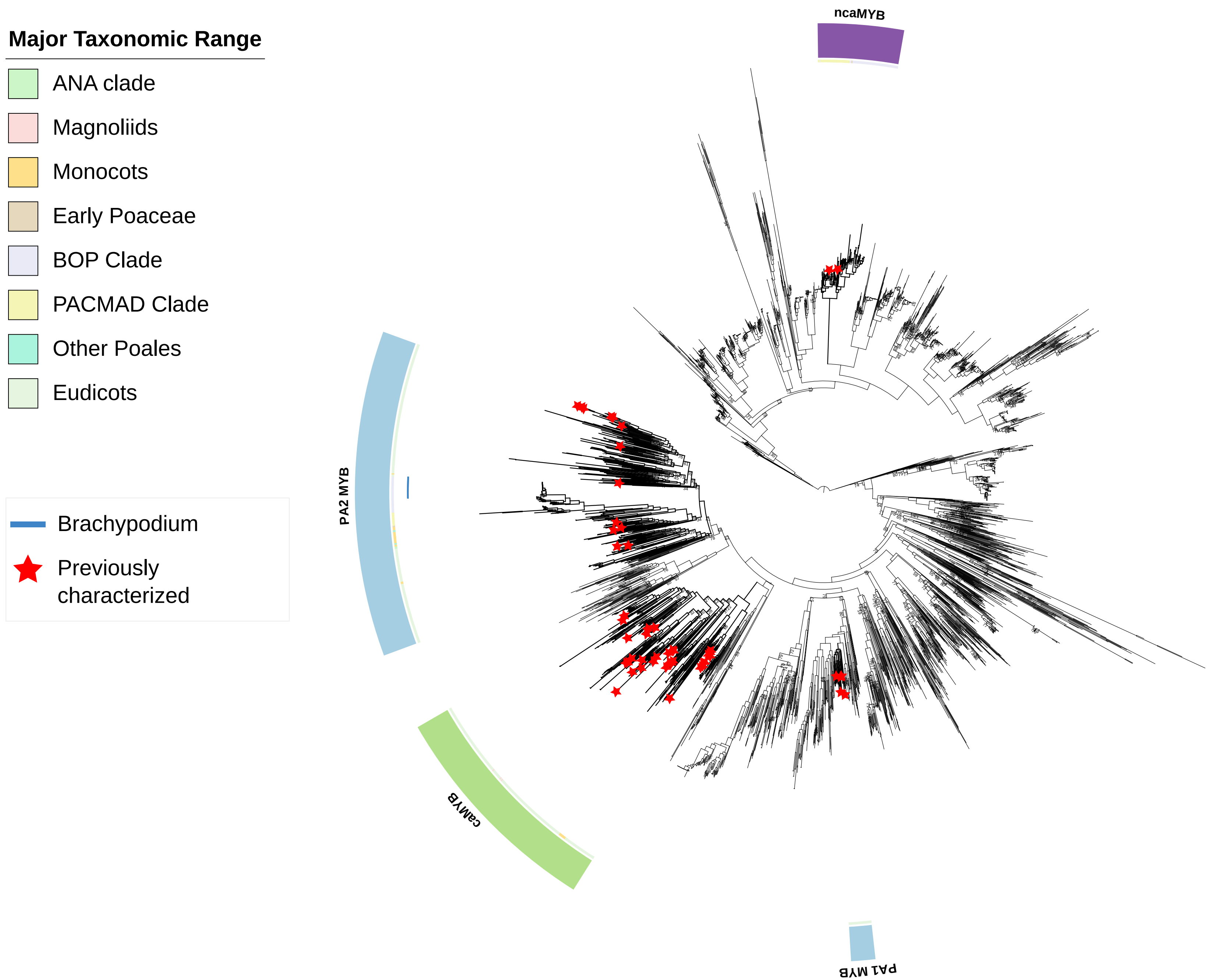


**Supplementary Figure 1:** Phylogenetic relationship of anthocyanin-related MYBs. The alignment is constructed using MAFFT G-INS-i method, followed by backtranslation to codons. Maximum-likelihood (ML) phylogenetic tree is constructed using IQ-TREE with GTR+I+R10 substitution model. Numbers above the nodes are the bootstrap values of ML analysis based on 1000 replicates. Bootstraps value <80% are shown. Leaf labels are hidden to reduce complexity.

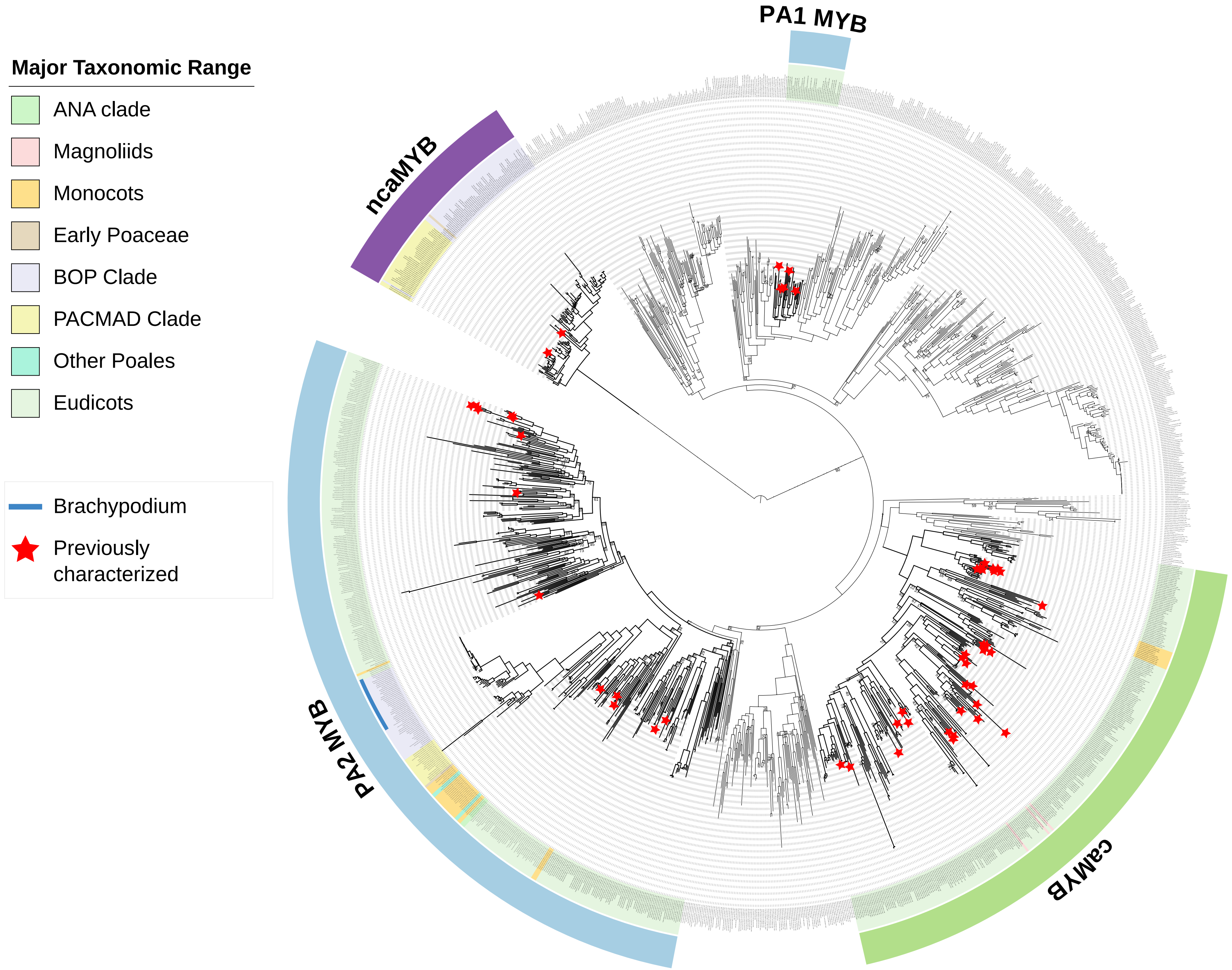


**Supplementary Figure 2:** Phylogenetic relationship of anthocyanin-related MYBs. This tree is pruned from the Supplementary Figure 1 tree to highlight the anthocyanin-related MYB clades. The alignment is constructed using MAFFT G-INS-i method, followed by backtranslation to codons. Maximum-likelihood (ML) phylogenetic tree is constructed using IQTREE with GTR+I+R10 substitution model. Numbers above the nodes are the bootstrap values of ML analysis based on 1000 replicates. Bootstraps value <80% are shown.

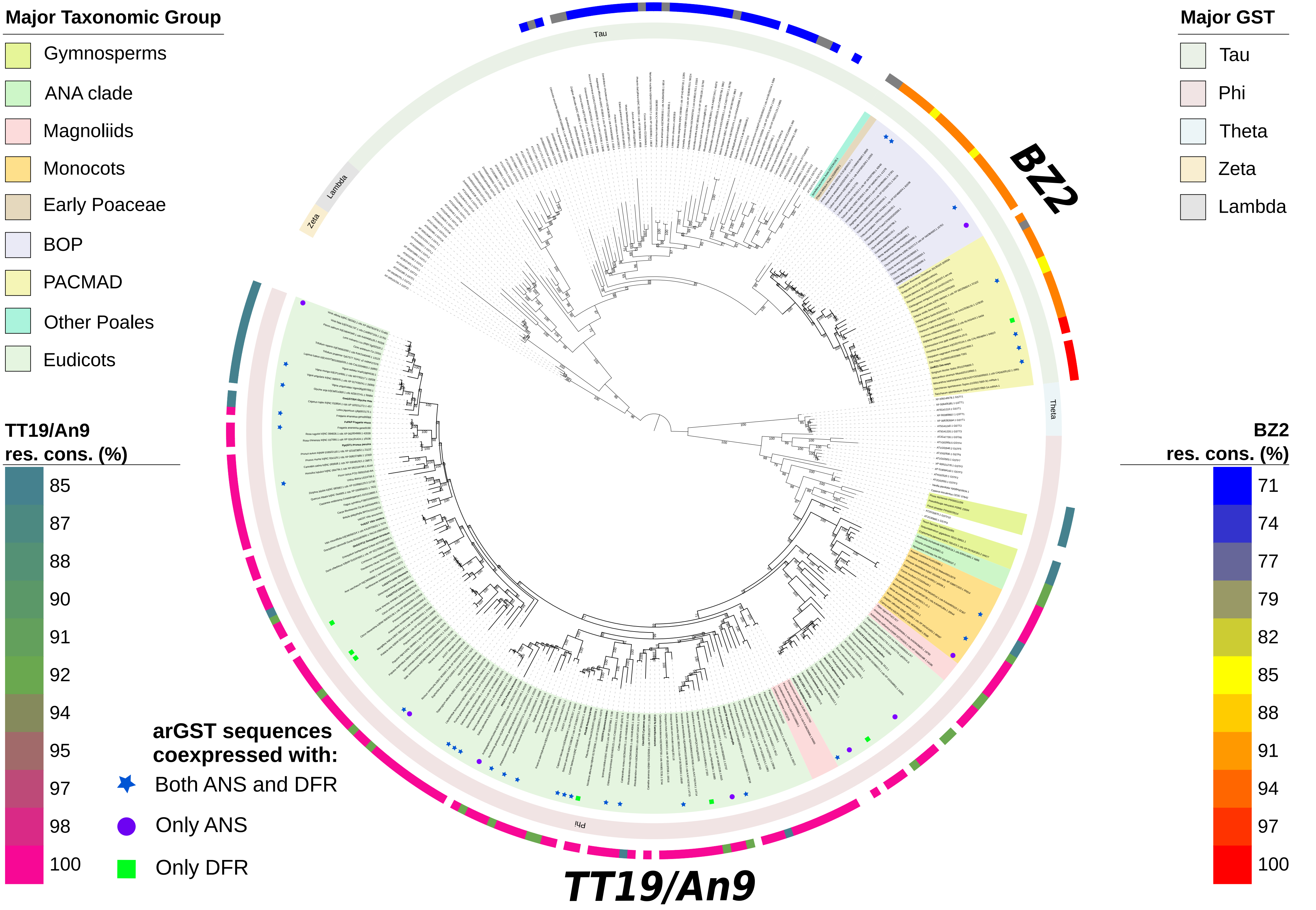


**Supplementary Figure 3:** Phylogenetic relationship of arGSTs. The functional arGST sequences were identified using KIPEs3 and coexpression analysis. The alignment is constructed using MAFFT G-INS-i method, followed by backtranslation to codon. Maximum-likelihood (ML) phylogenetic tree is constructed using IQTREE with SYM+R7 substitution model. Numbers above the nodes are the bootstrap values of ML analysis based on 1000 replicates.

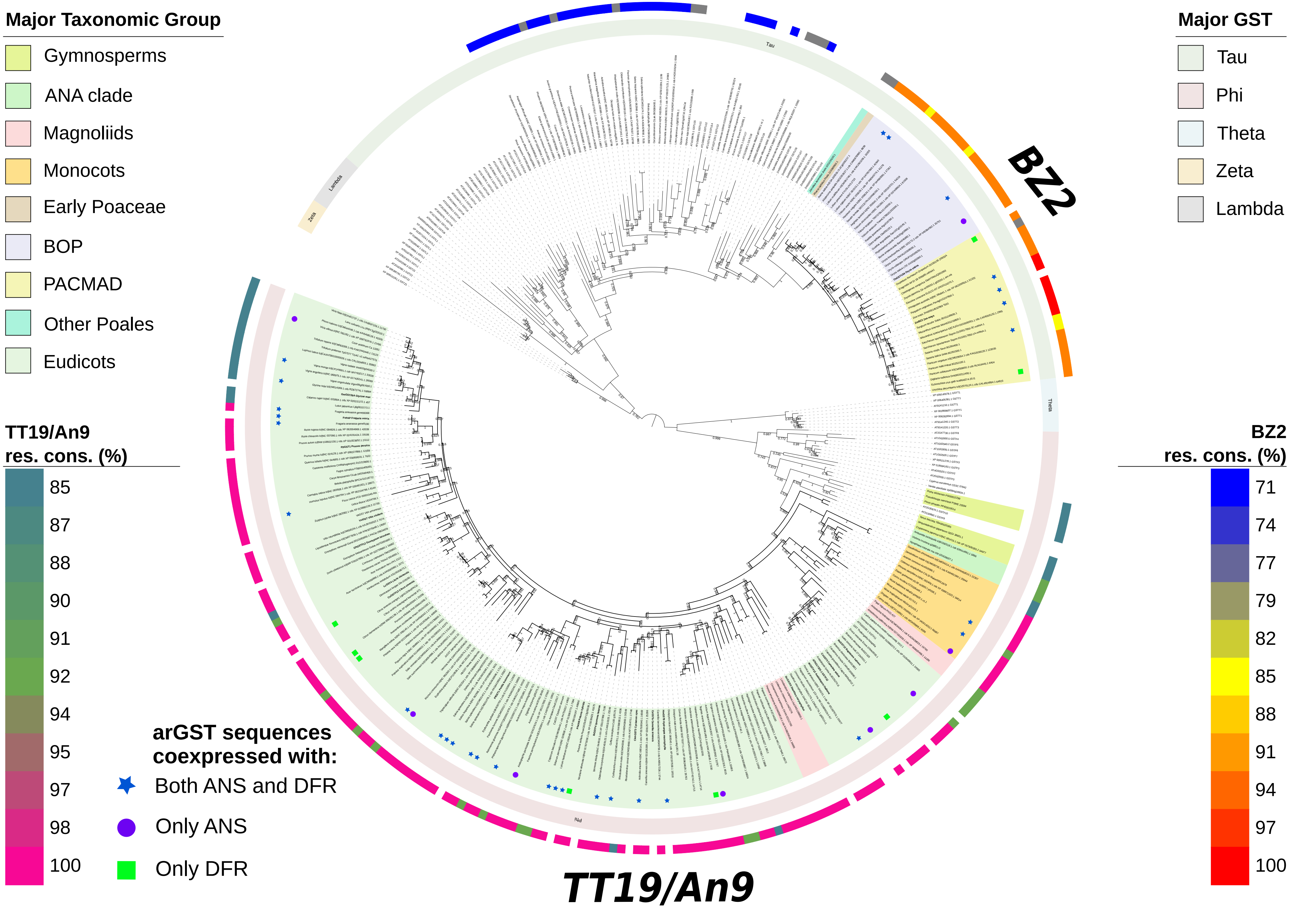


**Supplementary Figure 4:** Phylogenetic relationship of arGSTs. The functional arGST sequences were identified using KIPEs3 and coexpression analysis. The alignment is constructed using MAFFT G-INS-i method, followed by backtranslation to codon. Maximum-likelihood (ML) phylogenetic tree is constructed using FastTree with Jukes-Cantor + CAT model. Numbers above the nodes are the bootstrap values of ML analysis based on 1000 replicates (0-1 based).

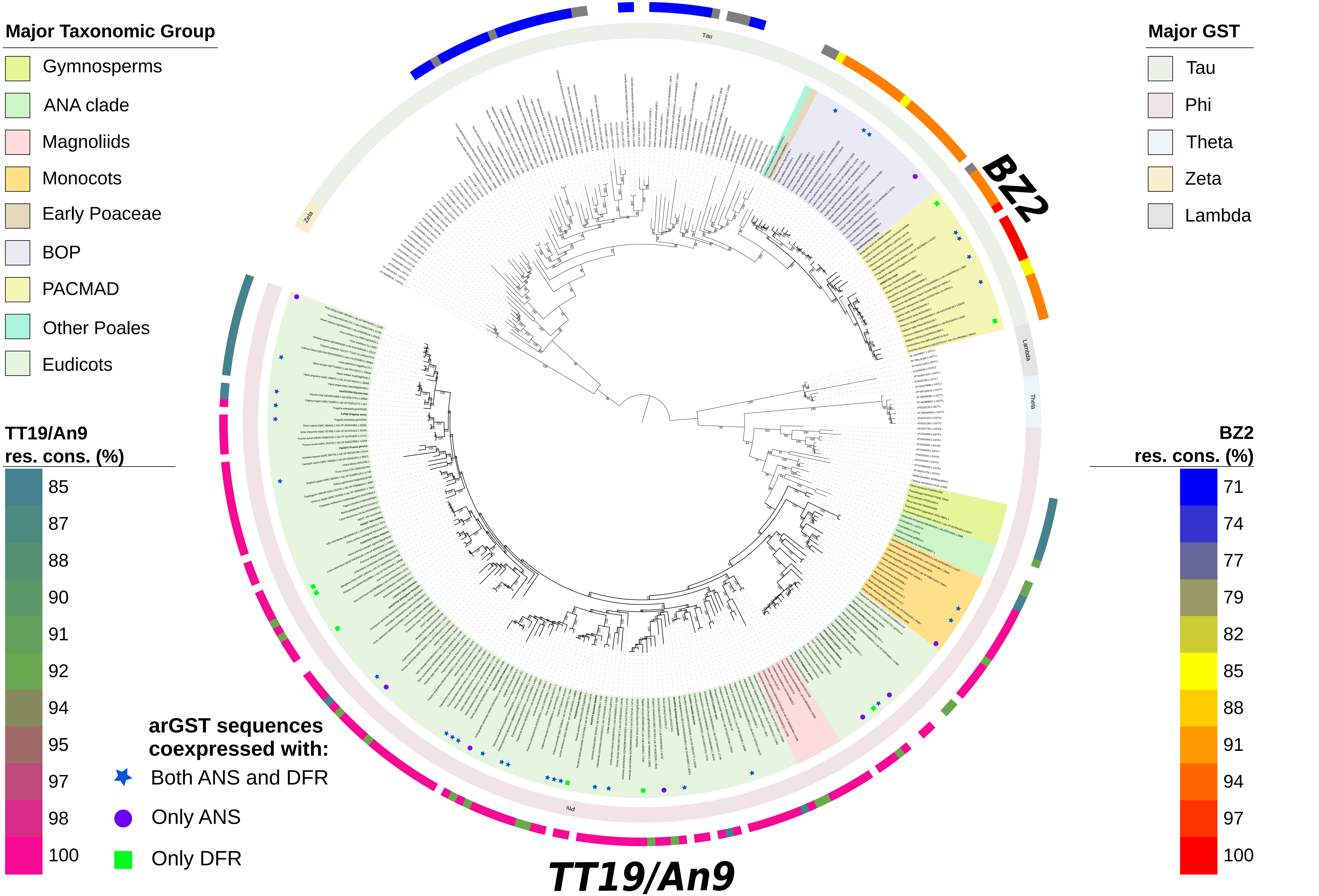


**Supplementary Figure 5:** Phylogenetic relationship of arGSTs. The functional arGST sequences were identified using KIPEs3 and coexpression analysis. The alignment is constructed using MUSCLE, followed by backtranslation to codon. Maximum-likelihood (ML) phylogenetic tree is constructed using IQTREE with TVMe+I+R7 substitution model. Numbers above the nodes are the bootstrap values of ML analysis based on 1000 replicates.

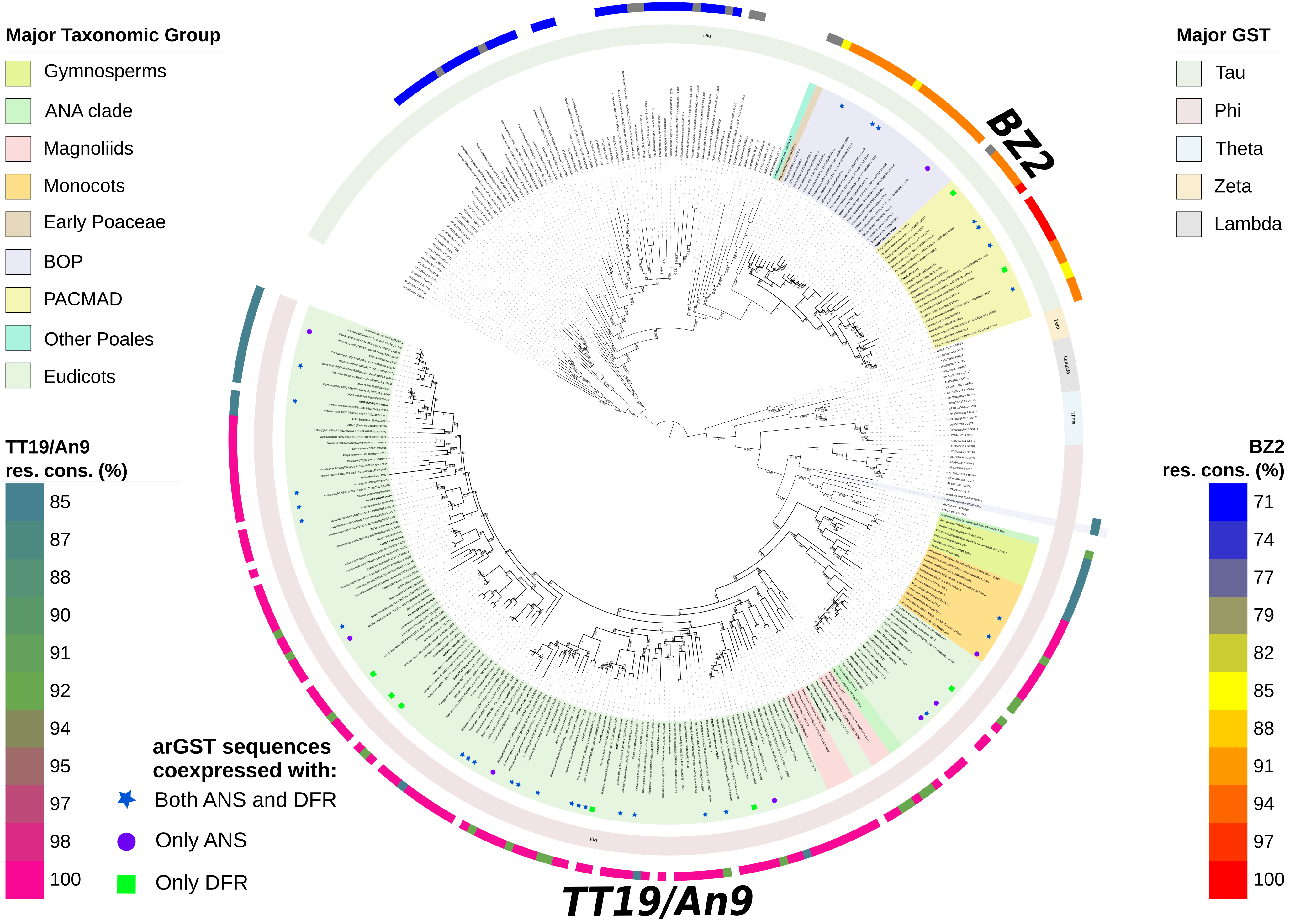


**Supplementary Figure 6:** Phylogenetic relationship of arGSTs. The functional arGST sequences were identified using KIPEs3 and coexpression analysis. The alignment is constructed using MUSCLE, followed by backtranslation to codon. Maximum-likelihood (ML) phylogenetic tree is constructed using FastTree with Jukes-Cantor + CAT model. Numbers above the nodes are the bootstrap values of ML analysis based on 1000 replicates (0-1 based).

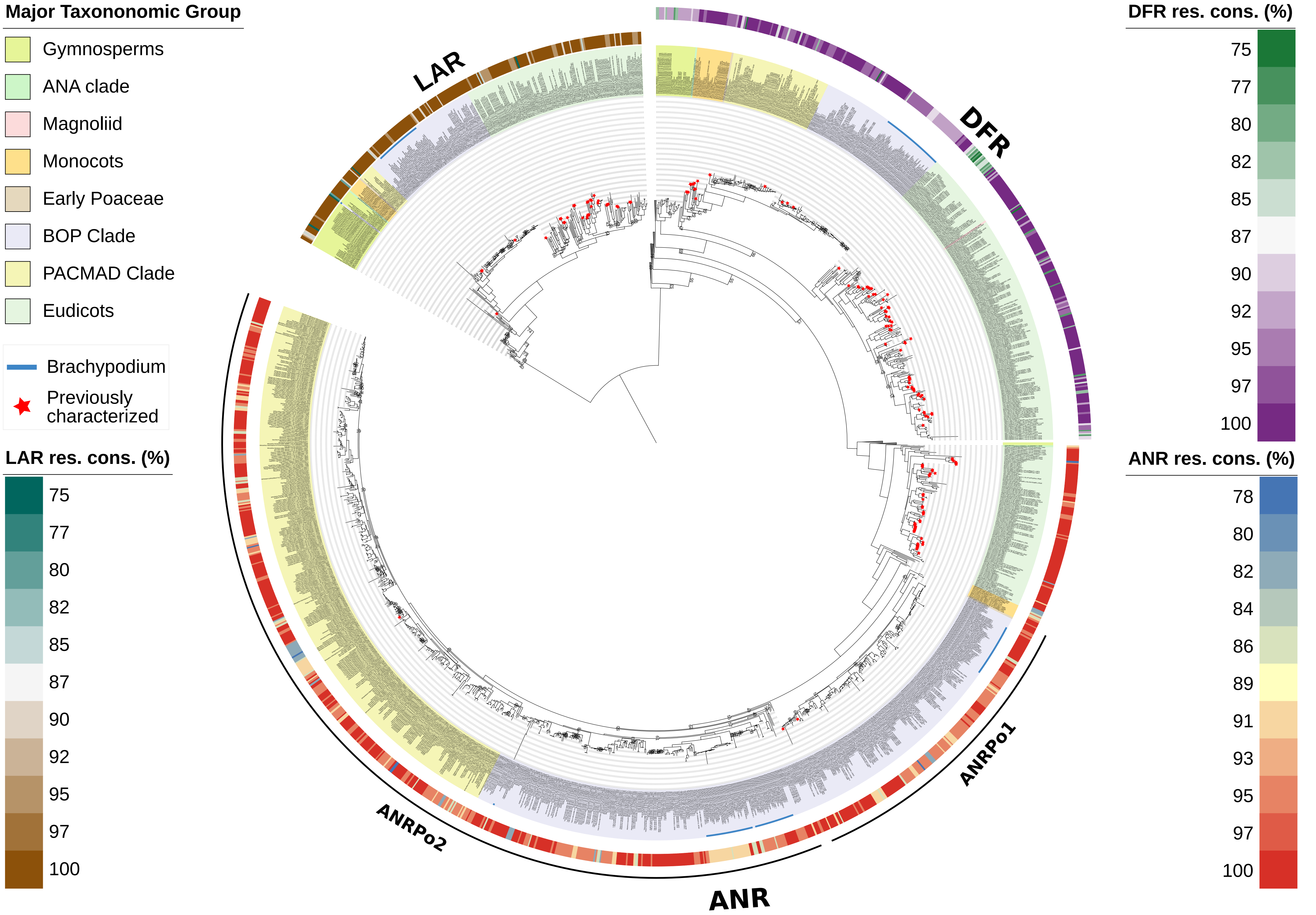


**Supplementary Figure 7:** Phylogenetic relationship of SDR family members participating flavonoid biosynthesis (LAR, DFR, and ANR). The alignment is constructed using MAFFT G-INS-i method, followed by backtranslation to codon. Maximum-likelihood (ML) phylogenetic tree is constructed using IQ-TREE with GTR+ASC+R10 substitution model. Numbers above the nodes are the bootstrap values of ML analysis based on 1000 replicates.

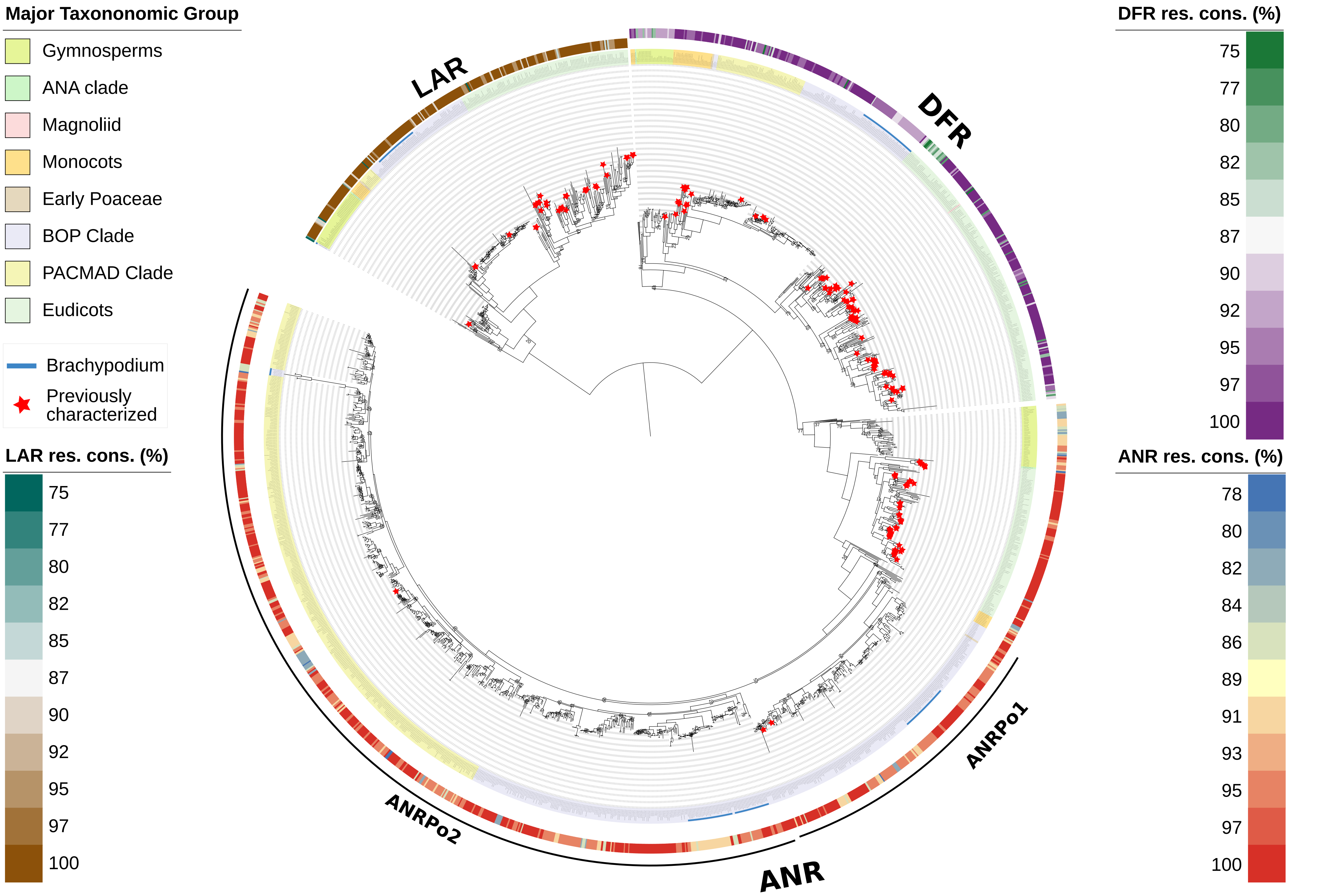


**Supplementary Figure 8:** Phylogenetic relationship of SDR family members participating flavonoid biosynthesis (LAR, DFR, and ANR). The alignment is constructed using MAFFT G-INS-i method, followed by backtranslation to codon. Maximum-likelihood (ML) phylogenetic tree is constructed using FastTree with Jukes-Cantor + CAT model. Numbers above the nodes are the bootstrap values of ML analysis based on 1000 replicates. Bootstrap values <80% are shown.

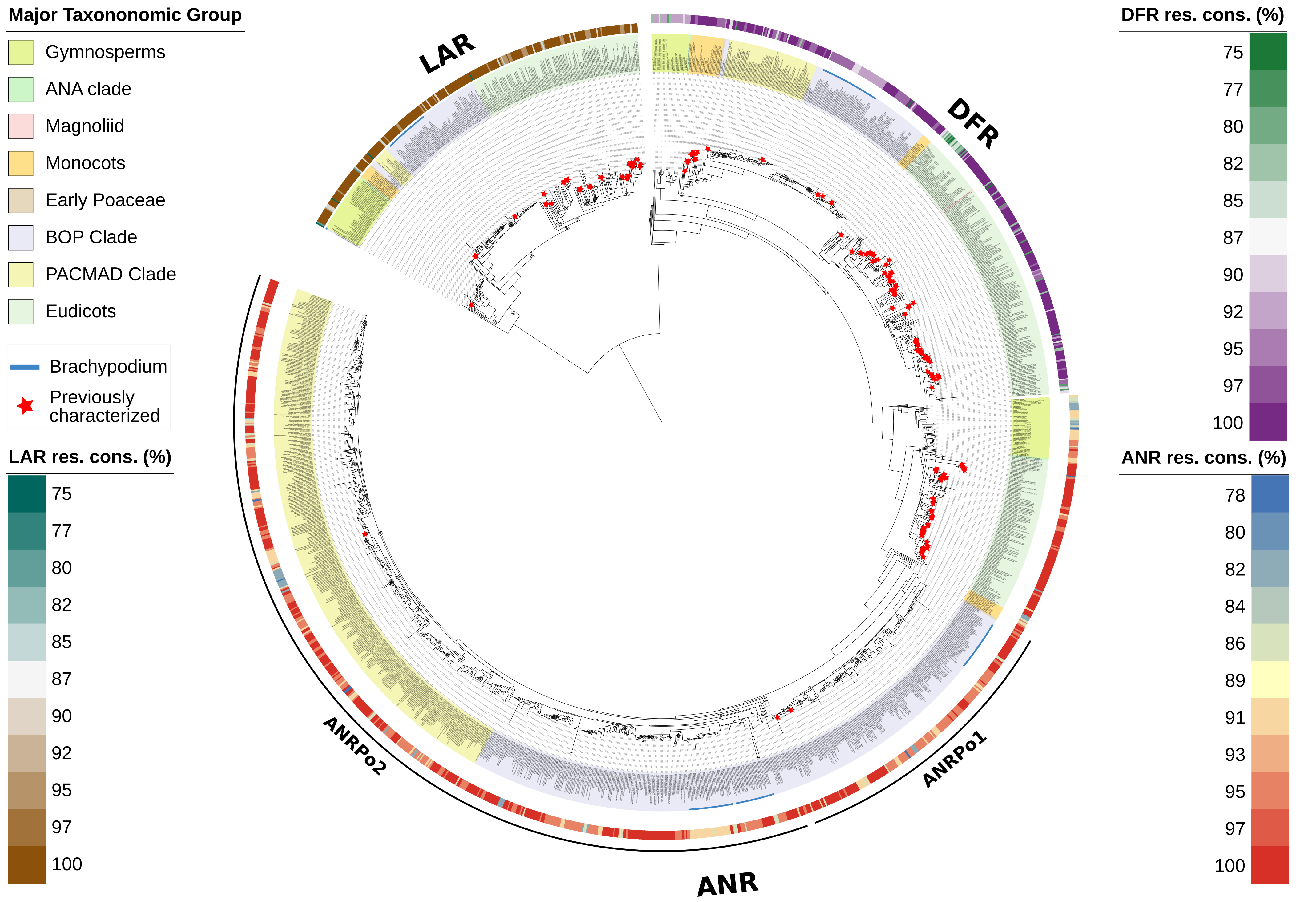


**Supplementary Figure 9:** Phylogenetic relationship of SDR family members participating flavonoid biosynthesis (LAR, DFR, and ANR). The alignment is constructed using MUSCLE, followed by backtranslation to codon. Maximum-likelihood (ML) phylogenetic tree is constructed using IQ-TREE with GTR+ASC+R10 substitution model. Numbers above the nodes are the bootstrap values of ML analysis based on 1000 replicates. Bootstrap values <80% are shown.

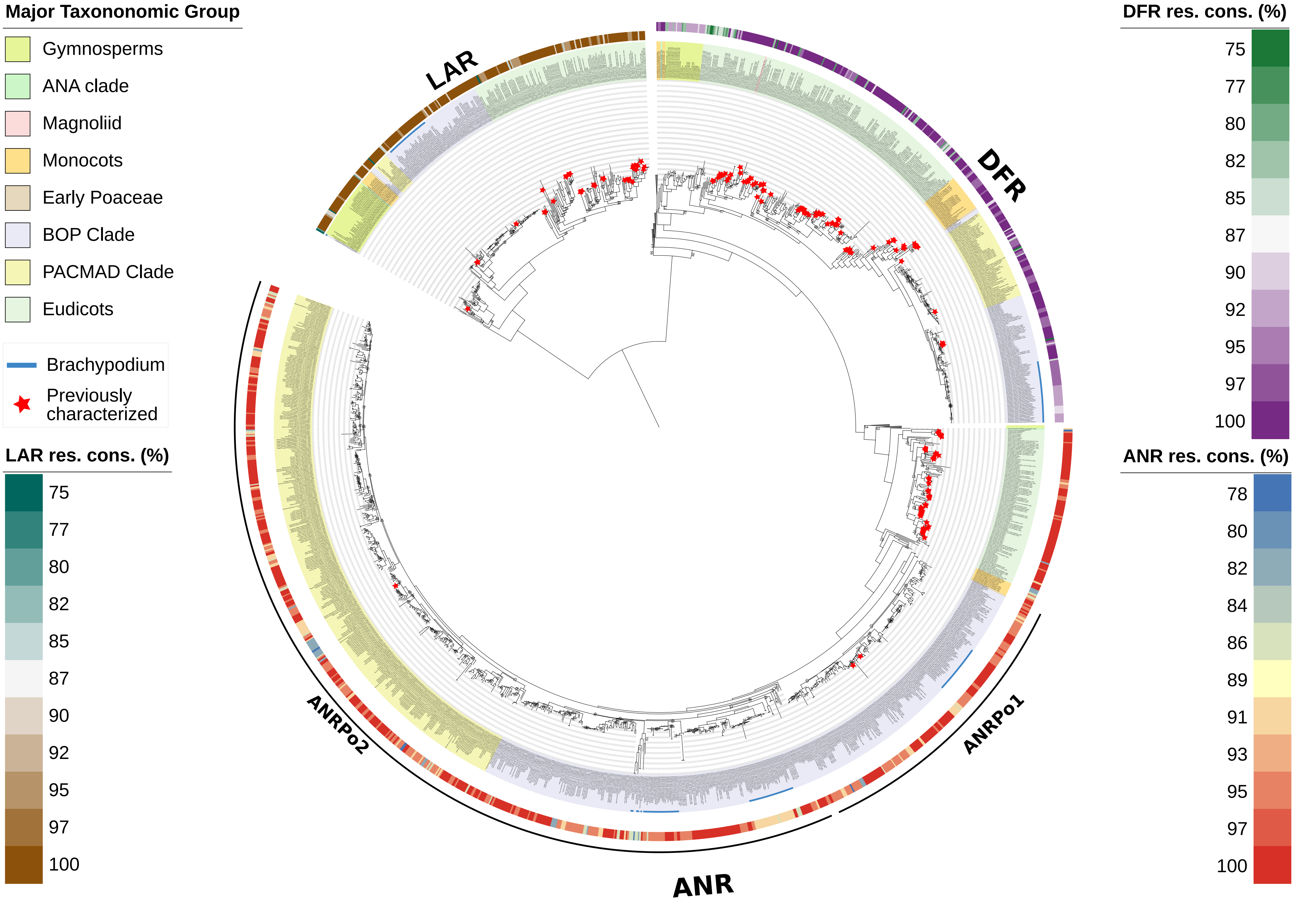


**Supplementary Figure 10:** Phylogenetic relationship of SDR family members participating flavonoid biosynthesis (LAR, DFR, and ANR). The alignment is constructed using MUSCLE, followed by backtranslation to codon. Maximum-likelihood (ML) phylogenetic tree is constructed using FastTree with Jukes-Cantor + CAT model. Numbers above the nodes are the bootstrap values of ML analysis based on 1000 replicates. Bootstrap values <80% are shown.

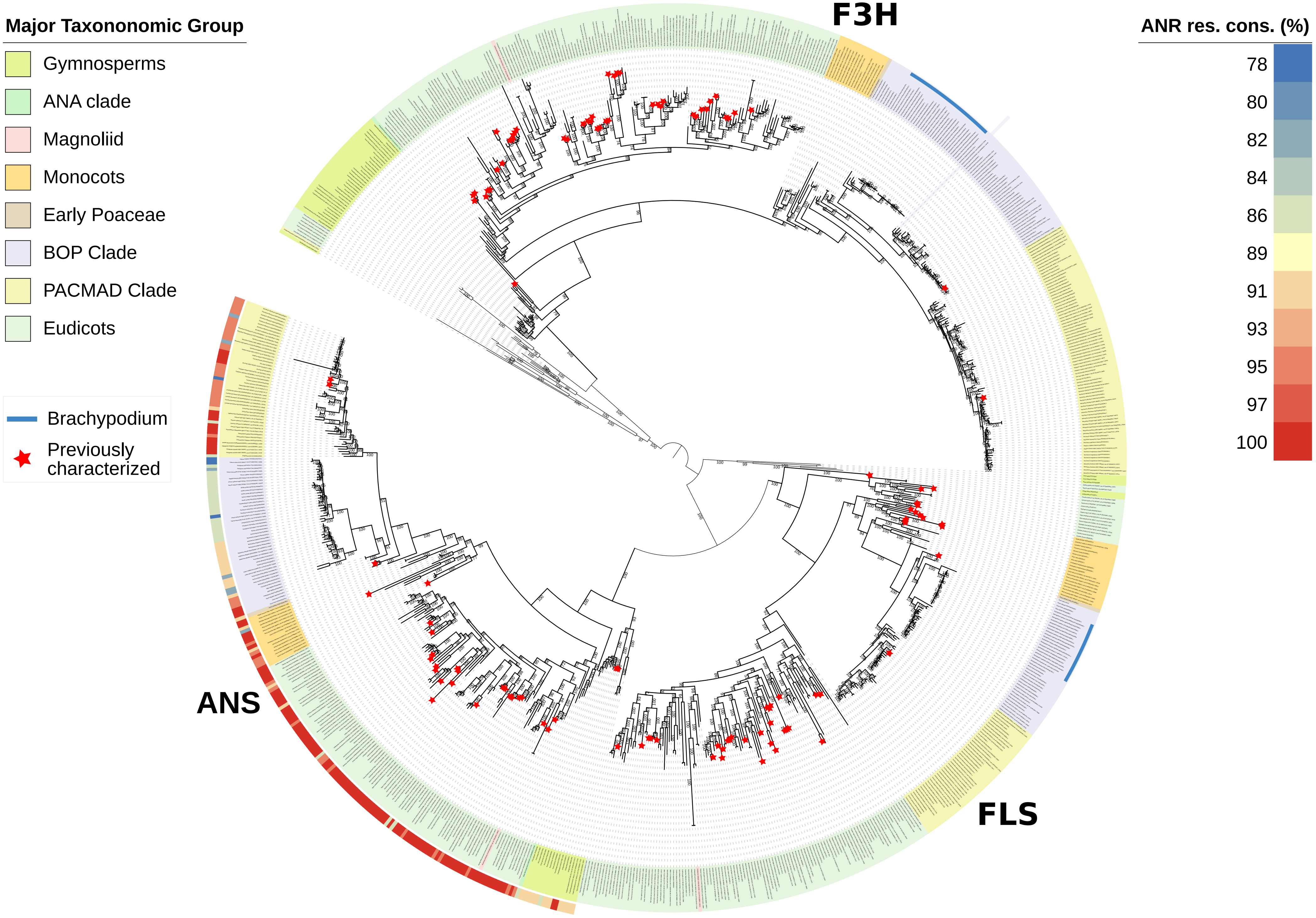


**Supplementary Figure 11:** Phylogenetic relationship of 2ODD family members participating in flavonoid biosynthesis (F3H, FLS, and ANS). The alignment is constructed using MAFFT using G-INS-i method, followed by backtranslation to codon. Maximum-likelihood (ML) phylogenetic tree is constructed using IQ-TREE with GTR+F+I+R10 substitution model. Numbers above the nodes are the bootstrap values of ML analysis based on 1000 replicates.

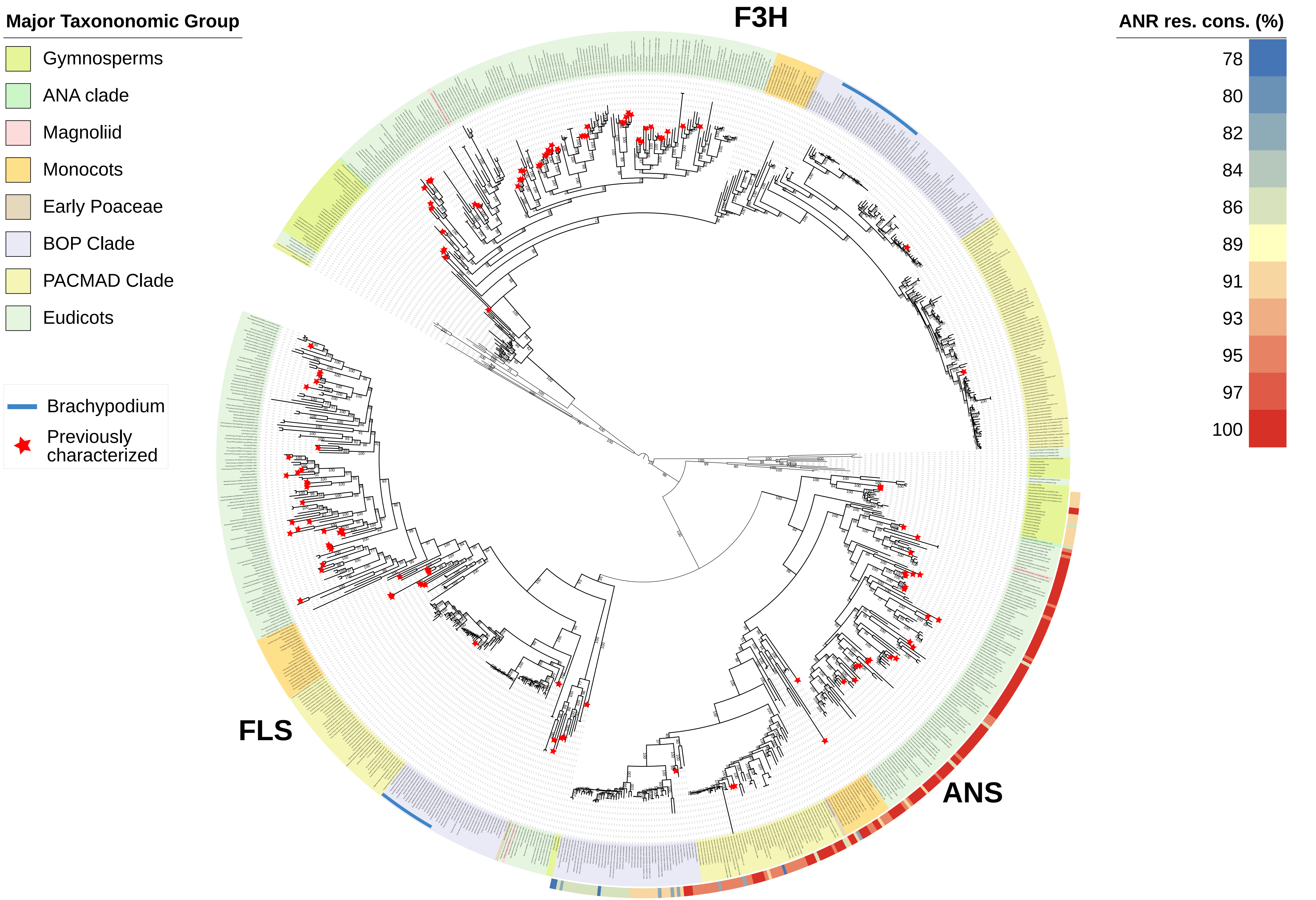


**Supplementary Figure 12:** Phylogenetic relationship of 2ODD family members participating in flavonoid biosynthesis (F3H, FLS, and ANS). The alignment is constructed using MAFFT using G-INS-i method, followed by backtranslation to codon. Maximum-likelihood (ML) phylogenetic tree is constructed using FastTree with Jukes-Cantor + CAT model. Numbers above the nodes are the bootstrap values of ML analysis based on 1000 replicates.

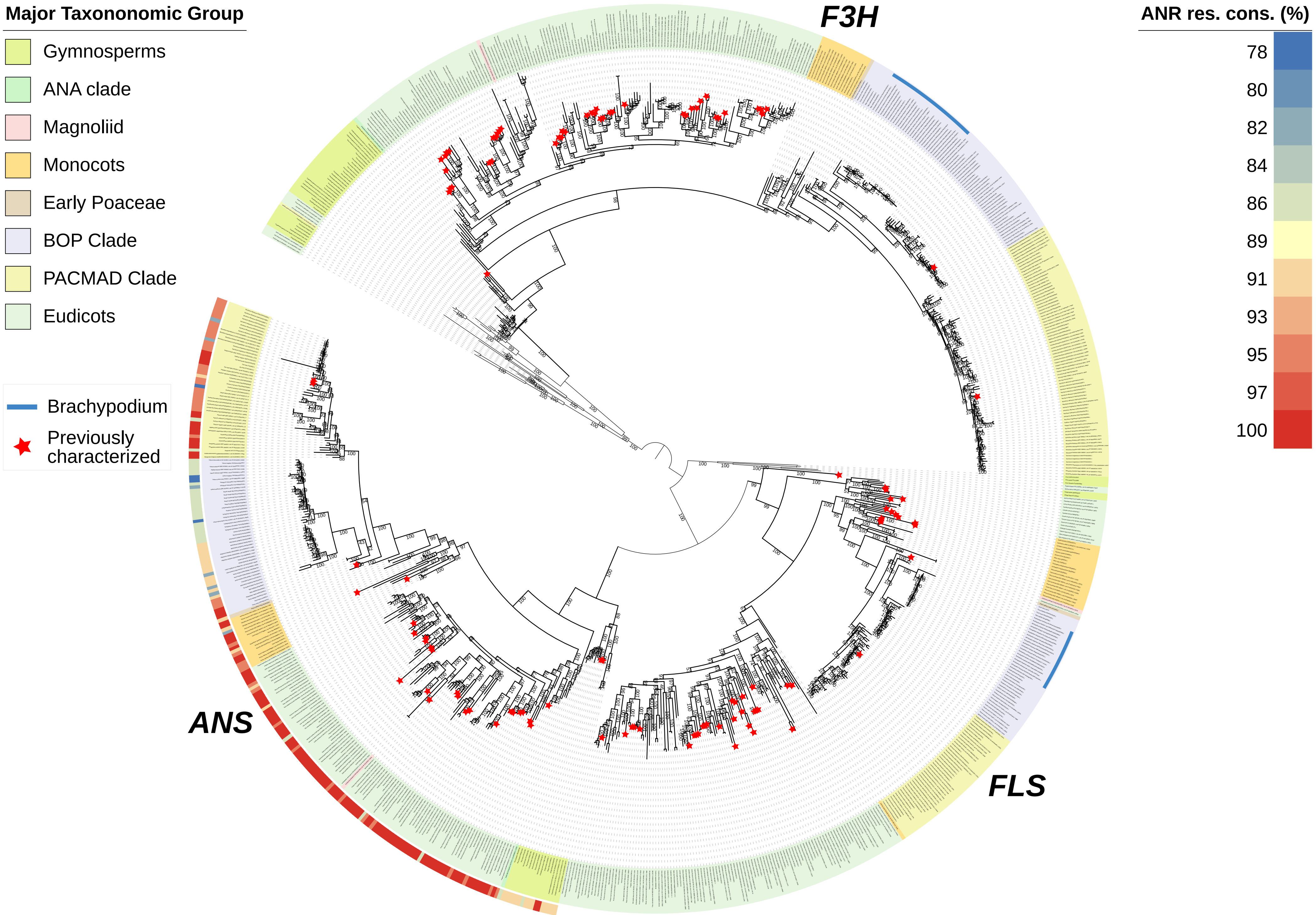


**Supplementary Figure 13:** Phylogenetic relationship of 2ODD family members participating in flavonoid biosynthesis (F3H, FLS, and ANS). The alignment is constructed using MUSCLE, followed by backtranslation to codon. Maximum-likelihood (ML) phylogenetic tree is constructed using IQ-TREE with GTR+F+I+R9 substitution model. Numbers above the nodes are the bootstrap values of ML analysis based on 1000 replicates.

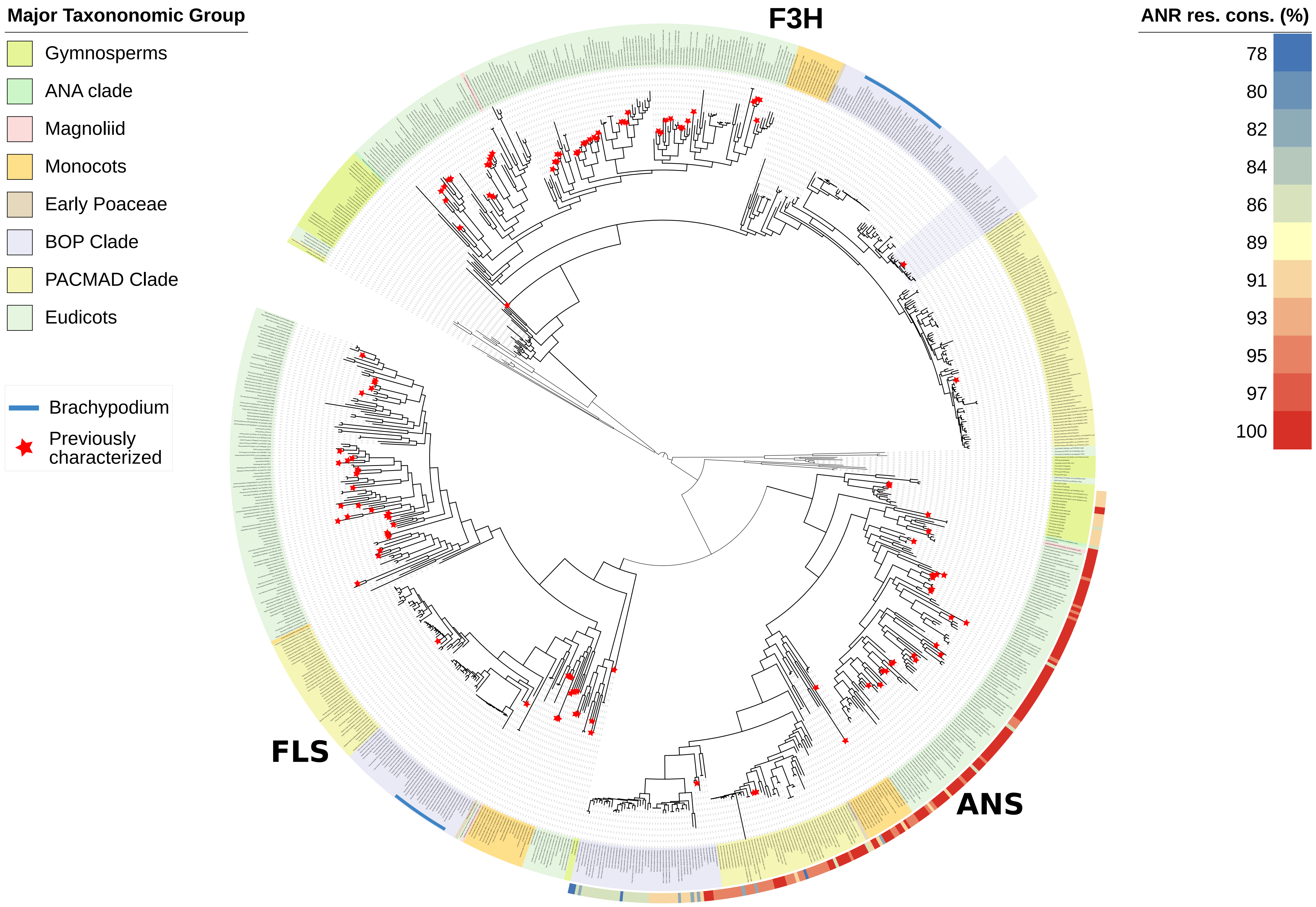


**Supplementary Figure 14:** Phylogenetic relationship of 2ODD family members participating in flavonoid biosynthesis (F3H, FLS, and ANS). The alignment is constructed using MUSCLE, followed by backtranslation to codon. Maximum-likelihood (ML) phylogenetic tree is constructed using FastTree with Jukes-Cantor + CAT model. Numbers above the nodes are the bootstrap values of ML analysis based on 1000 replicates.

Lovell J.T., MacQueen A.H., Mamidi S., Bonnette J., Jenkins J., Napier J.D., Sreedasyam A., Healey A., Session A., Shu S., Barry K., Bonos S., Boston L., Daum C., Deshpande S., Ewing A., Grabowski P.P., Haque T., Harrison M., Jiang J., Kudrna D., Lipzen A., Pendergast T.H., Plott C., Qi P., Saski C.A., Shakirov E.V., Sims D., Sharma M., Sharma R., Stewart A., Singan V.R., Tang Y., Thibivillier S., Webber J., Weng X., Williams M., Wu G.A., Yoshinaga Y., Zane M., Zhang L., Zhang J., Behrman K.D., Boe A.R., Fay P.A., Fritschi F.B., Jastrow J.D., Lloyd-Reilley J., Martínez-Reyna J.M., Matamala R., Mitchell R.B., Rouquette F.M., Ronald P., Saha M., Tobias C.M., Udvardi M., Wing R.A., Wu Y., Bartley L.E., Casler M., Devos K.M., Lowry D.B., Rokhsar D.S., Grimwood J., Juenger T.E., Schmutz

J. (2021) Genomic mechanisms of climate adaptation in polyploid bioenergy switchgrass. Nature **590**:438–444.

Luo M.-C., Gu Y.Q., Puiu D., Wang H., Twardziok S.O., Deal K.R., Huo N., Zhu T., Wang L., Wang Y., McGuire P.E., Liu S., Long H., Ramasamy R.K., Rodriguez J.C., Van S.L., Yuan L., Wang Z., Xia Z., Xiao L., Anderson O.D., Ouyang S., Liang Y., Zimin A.V., Pertea G., Qi P., Bennetzen J.L., Dai X., Dawson M.W., Müller H.-G., Kugler K., Rivarola-Duarte L., Spannagl M., Mayer K.F.X., Lu F.-H., Bevan M.W., Leroy P., Li P., You F.M., Sun Q., Liu Z., Lyons E., Wicker T., Salzberg S.L., Devos K.M., Dvořák J. (2017) Genome sequence of the progenitor of the wheat D genome Aegilops tauschii. Nature **551**:498–502.

Ma P.-F., Liu Y.-L., Jin G.-H., Liu J.-X., Wu H., He J., Guo Z.-H., Li D.-Z. (2021) The Pharus latifolius genome bridges the gap of early grass evolution. The Plant Cell **33**:846–864.

Ma L., Liu K.-W., Li Z., Hsiao Y.-Y., Qi Y., Fu T., Tang G.-D., Zhang D., Sun W.-H., Liu D.-K., Li Y., Chen G.-Z., Liu X.-D., Liao X.-Y., Jiang Y.-T., Yu X., Hao Y., Huang J., Zhao X.-W., Ke S., Chen Y.-Y., Wu W.-L., Hsu J.-L., Lin Y.-F., Huang M.-D., Li C.-Y., Huang L., Wang Z.-W., Zhao X., Zhong W.-Y., Peng D.-H., Ahmad S., Lan S., Zhang J.-S., Tsai W.-C., Van de Peer Y., Liu Z.-J. (2023) Diploid and tetraploid genomes of Acorus and the evolution of monocots. Nature Communications **14**:3661.

Ma X., Ru D., Morales-Briones D.F., Mei F., Wu J., Liu J., Wu S. (2023) Genome sequence and salinity adaptation of the desert shrub Nitraria sibirica (Nitrariaceae, Sapindales). DNA Research **30**:dsad011.

Mamidi S., Healey A., Huang P., Grimwood J., Jenkins J., Barry K., Sreedasyam A., Shu S., Lovell J.T., Feldman M., Wu J., Yu Y., Chen C., Johnson J., Sakakibara H., Kiba T., Sakurai T., Tavares R., Nusinow D.A., Baxter I., Schmutz J., Brutnell T.P., Kellogg E.A. (2020) A genome resource for green millet Setaria viridis enables discovery of agronomically valuable loci. Nature Biotechnology **38**:1203–1210.

Mao W., Yao G., Wang S., Zhou L., Chen G., Dong N., Hu G., Mao W., Yao G., Wang S., Zhou L., Chen G., Dong N., Hu G. (2021) Chromosome-level genomes of seeded and seedless date plum based on third-generation DNA sequencing and Hi-C analysis. Forestry Research **1** [online] URL: https://www.maxapress.com/article/doi/10.48130/FR-2021-0009 (accessed 21 September 2025).
Martinez-Hernandez J.E., Salvo-Garrido H., Levicoy D., Caligari P.D.S., Rupayán A., Moyano T., Carrasco M., Hernandez S., Armijo-Godoy G., Westermeyer F., Larama G. (2024) Chromosome-level genome assembly of yellow lupin (Lupinus luteus) provides novel insights into genome evolution, crop adaptation and seed protein in the three most cultivated lupins. [online] URL: https://www.researchsquare.com/article/rs-4171664/v1 (accessed 21 September 2025).

Ming R., VanBuren R., Wai C.M., Tang H., Schatz M.C., Bowers J.E., Lyons E., Wang M.-L., Chen J., Biggers E., Zhang J., Huang L., Zhang L., Miao W., Zhang J., Ye Z., Miao C., Lin Z., Wang H., Zhou H., Yim W.C., Priest H.D., Zheng C., Woodhouse M., Edger P.P., Guyot R., Guo H.-B., Guo H., Zheng G., Singh R., Sharma A., Min X., Zheng Y., Lee H., Gurtowski J., Sedlazeck F.J., Harkess A., McKain M.R., Liao Z., Fang J., Liu J., Zhang X., Zhang Q., Hu W., Qin Y., Wang K., Chen L.-Y., Shirley N., Lin Y.-R., Liu L.-Y., Hernandez A.G., Wright C.L., Bulone V., Tuskan G.A., Heath K., Zee F., Moore P.H., Sunkar R., Leebens-Mack J.H., Mockler T., Bennetzen J.L., Freeling M., Sankoff D., Paterson

A.H., Zhu X., Yang X., Smith J.A.C., Cushman J.C., Paull R.E., Yu Q. (2015) The pineapple genome and the evolution of CAM photosynthesis. Nature Genetics **47**:1435–1442.

Minoji K., Sakai T. (2024) A chromosome-scale genome assembly of Timorese crabgrass (Digitaria radicosa): a useful genomic resource for the Poaceae. G3 Genes|Genomes|Genetics **14**:jkae242.
Nowak M.S., Harder B., Meckoni S.N., Friedhoff R., Wolff K., Pucker B. (2025) Genome sequence and RNA-seq analysis reveal genetic basis of flower coloration in the giant water lily Victoria cruziana. :2024.06.15.599162. [online] URL: https://www.biorxiv.org/content/10.1101/2024.06.15.599162v3 (accessed 21 September 2025).

Paterson A.H., Bowers J.E., Bruggmann R., Dubchak I., Grimwood J., Gundlach H., Haberer G., Hellsten U., Mitros T., Poliakov A., Schmutz J., Spannagl M., Tang H., Wang X., Wicker T., Bharti A.K., Chapman J., Feltus F.A., Gowik U., Grigoriev I.V., Lyons E., Maher C.A., Martis M., Narechania A., Otillar R.P., Penning B.W., Salamov A.A., Wang Y., Zhang L., Carpita N.C., Freeling M., Gingle A.R., Hash C.T., Keller B., Klein P., Kresovich S., McCann M.C., Ming R., Peterson D.G., Mehboob-ur-Rahman, Ware D., Westhoff P., Mayer K.F.X., Messing J., Rokhsar D.S. (2009) The Sorghum bicolor genome and the diversification of grasses. Nature **457**:551–556.

Pereira L., Alenazi A.S., Mian S., Leitch I.J., Christin P.-A., Osborne C.P., Dunning L.T. (2025) Gene turnover and contingency facilitated the repeated evolution of C4 photosynthesis in grasses. :2025.04.23.650007. [online] URL: https://www.biorxiv.org/content/10.1101/2025.04.23.650007v1 (accessed 21 September 2025).
Phtheirospermum japonicum genome assembly Pjver1 NCBI [online] URL: https://www.ncbi.nlm.nih.gov/datasets/genome/GCA_014905375.1/ (accessed 21 September 2025).
P.latifolius v1.1: Phytozome [online] URL: https://phytozome-next.jgi.doe.gov/info/Platifolius_v1_1 (accessed 21 September 2025).

Qin L., Hu Y., Wang J., Wang X., Zhao R., Shan H., Li K., Xu P., Wu H., Yan X., Liu L., Yi X., Wanke S., Bowers J.E., Leebens-Mack J.H., dePamphilis C.W., Soltis P.S., Soltis D.E., Kong H., Jiao Y. (2021) Insights into angiosperm evolution, floral development and chemical biosynthesis from the Aristolochia fimbriata genome. Nature Plants **7**:1239–1253.

Rabanus-Wallace M.T., Hackauf B., Mascher M., Lux T., Wicker T., Gundlach H., Baez M., Houben A., Mayer K.F.X., Guo L., Poland J., Pozniak C.J., Walkowiak S., Melonek J., Praz C.R., Schreiber M., Budak H., Heuberger M., Steuernagel B., Wulff B., Börner A., Byrns B., Čížková J., Fowler D.B., Fritz A., Himmelbach A., Kaithakottil G., Keilwagen J., Keller B., Konkin D., Larsen J., Li Q., Myśków B., Padmarasu S., Rawat N., Sesiz U., Biyiklioglu-Kaya S., Sharpe A., Šimková H., Small I., Swarbreck D., Toegelová H., Tsvetkova N., Voylokov A.V., Vrána J., Bauer E., Bolibok-Bragoszewska H., Doležel J., Hall A., Jia J., Korzun V., Laroche A., Ma X.-F., Ordon F., Özkan H., Rakoczy-Trojanowska M., Scholz U., Schulman A.H., Siekmann D., Stojałowski S., Tiwari V.K., Spannagl M., Stein N. (2021) Chromosome-scale genome assembly provides insights into rye biology, evolution and agronomic potential. Nature Genetics **53**:564–573.

Revolinski S.R., Maughan P.J., Coleman C.E., Burke I.C. (2023) Preadapted to adapt: underpinnings of adaptive plasticity revealed by the downy brome genome. Communications Biology **6**:326.

Rhododendron molle genome assembly RHMOLv1 NCBI [online] URL: https://www.ncbi.nlm.nih.gov/datasets/genome/GCA_025413875.1/ (accessed 21 September 2025).

Ryan C., Fraser F., Irish N., Barker T., Knitlhoffer V., Durrant A., Reynolds G., Kaithakottil G., Swarbreck D., De Vega J.J. (2025) A haplotype-resolved chromosome-level genome assembly of Urochloa decumbens cv. Basilisk resolves its allopolyploid ancestry and composition. G3 Genes|Genomes|Genetics **15**:jkaf005.

Scott A.D., Zimin A.V., Puiu D., Workman R., Britton M., Zaman S., Caballero M., Read A.C., Bogdanove A.J., Burns E., Wegrzyn J., Timp W., Salzberg S.L., Neale D.B. (2020) A Reference Genome Sequence for Giant Sequoia. G3 Genes|Genomes|Genetics **10**:3907–3919.

Studer A.J., Schnable J.C., Weissmann S., Kolbe A.R., McKain M.R., Shao Y., Cousins A.B., Kellogg E.A., Brutnell T.P. (2016) The draft genome of the C3 panicoid grass species Dichanthelium oligosanthes. Genome Biology **17**:223.

Sun G., Wase N., Shu S., Jenkins J., Zhou B., Torres-Rodríguez J.V., Chen C., Sandor L., Plott C., Yoshinga Y., Daum C., Qi P., Barry K., Lipzen A., Berry L., Pedersen C., Gottilla T., Foltz A., Yu H., O’Malley R., Zhang C., Devos K.M., Sigmon B., Yu B., Obata T., Schmutz J., Schnable J.C. (2022) Genome of Paspalum vaginatum and the role of trehalose mediated autophagy in increasing maize biomass. Nature Communications **13**:7731.

T.intermedium v3.1: Phytozome [online] URL: https://phytozome-next.jgi.doe.gov/info/Tintermedium_v3_1 (accessed 21 September 2025).

Vega J.M., Podio M., Orjuela J., Siena L.A., Pessino S.C., Combes M.C., Mariac C., Albertini E., Pupilli F., Ortiz J.P.A., Leblanc O. (2024) Chromosome-scale genome assembly and annotation of Paspalum notatum Flüggé var. saurae. Scientific Data **11**:891.

Vignale F.A., Hernandez Garcia A., Modenutti C.P., Sosa E.J., Defelipe L.A., Oliveira R., Nunes G.L., Acevedo R.M., Burguener G.F., Rossi S.M., Zapata P.D., Marti D.A., Sansberro P., Oliveira G., Catania E.M., Smith M.N., Dubs N.M., Nair S., Barkman T.J., Turjanski A.G. (2025) Yerba mate (Ilex paraguariensis) genome provides new insights into convergent evolution of caffeine biosynthesis (D. Weigel, Ed.). eLife **14**:e104759.

Vogel J.P., Garvin D.F., Mockler T.C., Schmutz J., Rokhsar D., Bevan M.W., Barry K., Lucas S., Harmon-Smith M., Lail K., Tice H., Schmutz (Leader) J., Grimwood J., McKenzie N., Bevan M.W., Huo N., Gu Y.Q., Lazo G.R., Anderson O.D., Vogel (Leader) J.P., You F.M., Luo M.-C., Dvorak J., Wright J., Febrer M., Bevan M.W., Idziak D., Hasterok R., Garvin D.F., Lindquist E., Wang M., Fox S.E., Priest H.D., Filichkin S.A., Givan S.A., Bryant D.W., Chang J.H., Mockler (Leader) T.C., Wu H., Wu W., Hsia A.-P., Schnable P.S., Kalyanaraman A., Barbazuk B., Michael T.P., Hazen S.P., Bragg J.N., Laudencia-Chingcuanco D., Vogel J.P., Garvin D.F., Weng Y., McKenzie N., Bevan M.W., Haberer G., Spannagl M., Mayer (Leader) K., Rattei T., Mitros T., Rokhsar D., Lee S.-J., Rose J.K.C., Mueller L.A., York T.L., Wicker (Leader) T., Buchmann J.P., Tanskanen J., Schulman (Leader) A.H., Gundlach H., Wright J., Bevan M., Costa de Oliveira A., da C. Maia L., Belknap W., Gu Y.Q., Jiang N., Lai J., Zhu L., Ma J., Sun C., Pritham E., Salse (Leader) J., Murat F., Abrouk M., Haberer G., Spannagl M.,

Mayer K., Bruggmann R., Messing J., You F.M., Luo M.-C., Dvorak J., Fahlgren N., Fox S.E., Sullivan C.M., Mockler T.C., Carrington J.C., Chapman E.J., May G.D., Zhai J., Ganssmann M., Guna Ranjan Gurazada S., German M., Meyers B.C., Green (Leader) P.J., Bragg J.N., Tyler L., Wu J., Gu Y.Q., Lazo G.R., Laudencia-Chingcuanco D., Thomson J., Vogel (Leader) J.P., Hazen S.P., Chen S., Scheller H.V., Harholt J., Ulvskov P., Fox S.E., Filichkin S.A., Fahlgren N., Kimbrel J.A., Chang J.H., Sullivan C.M., Chapman E.J., Carrington J.C., Mockler T.C., Bartley L.E., Cao P., Jung K.-H., Sharma M.K., Vega-Sanchez M., Ronald P., Dardick C.D., De Bodt S., Verelst W., Inzé D., Heese M., Schnittger A., Yang X., Kalluri U.C., Tuskan G.A., Hua Z., Vierstra R.D., Garvin D.F., Cui Y., Ouyang S., Sun Q., Liu Z., Yilmaz A., Grotewold E., Sibout R., Hematy K., Mouille G., Höfte H., Michael T., Pelloux J., O'Connor D., Schnable J., Rowe S., Harmon F., Cass C.L., Sedbrook J.C., Byrne M.E., Walsh S., Higgins J., Bevan M., Li P., Brutnell T., Unver T., Budak H., Belcram H., Charles M., Chalhoub B., Baxter I., The International Brachypodium Initiative, Principal investigators, DNA sequencing and assembly, Pseudomolecule assembly and BAC end sequencing, Transcriptome sequencing and analysis, Gene analysis and annotation, Repeats analysis, Comparative genomics, Small RNA analysis, Manual annotation and gene family analysis (2010) Genome sequencing and analysis of the model grass Brachypodium distachyon. Nature **463**:763–768.

Wang J., Tian S., Sun X., Cheng X., Duan N., Tao J., Shen G. (2020) Construction of Pseudomolecules for the Chinese Chestnut (Castanea mollissima) Genome. G3 Genes|Genomes|Genetics **10**:3565–3574.
Wickett N.J., Mirarab S., Nguyen N., Warnow T., Carpenter E., Matasci N., Ayyampalayam S., Barker M.S., Burleigh J.G., Gitzendanner M.A., Ruhfel B.R., Wafula E., Der J.P., Graham S.W., Mathews S., Melkonian M., Soltis D.E., Soltis P.S., Miles N.W., Rothfels C.J., Pokorny L., Shaw A.J., DeGironimo L., Stevenson D.W., Surek B., Villarreal J.C., Roure B., Philippe H., dePamphilis C.W., Chen T., Deyholos M.K., Baucom R.S., Kutchan T.M., Augustin M.M., Wang J., Zhang Y., Tian Z., Yan Z., Wu X., Sun X., Wong G.K.-S., Leebens-Mack J. (2014) Phylotranscriptomic analysis of the origin and early diversification of land plants. Proceedings of the National Academy of Sciences **111**:E4859–E4868.

Wright J., Baker K., Barker T., Catchpole L., Durrant A., Fraser F., Gharbi K., Harrison C., Henderson S., Irish N., Kaithakottil G., Leitch I.J., Li J., Lucchini S., Neve P., Powell R., Rees H., Swarbreck D., Watkins C., Wood J., McTaggart S., Hall A., MacGregor D. (2024) Chromosome-scale genome assembly and de novo annotation of Alopecurus aequalis. Scientific Data **11**:1368.

Wu S., Lau K.H., Cao Q., Hamilton J.P., Sun H., Zhou C., Eserman L., Gemenet D.C., Olukolu B.A., Wang H., Crisovan E., Godden G.T., Jiao C., Wang X., Kitavi M., Manrique-Carpintero N., Vaillancourt B., Wiegert-Rininger K., Yang X., Bao K., Schaff J., Kreuze J., Gruneberg W., Khan A., Ghislain M., Ma D., Jiang J., Mwanga R.O.M., Leebens-Mack J., Coin L.J.M., Yencho G.C., Buell C.R., Fei Z. (2018) Genome sequences of two diploid wild relatives of cultivated sweetpotato reveal targets for genetic improvement. Nature Communications **9**:4580.

Wu D., Shen E., Jiang B., Feng Y., Tang W., Lao S., Jia L., Lin H.-Y., Xie L., Weng X., Dong C., Qian Q., Lin F., Xu H., Lu H., Cutti L., Chen H., Deng S., Guo L., Chuah T.-S., Song B.-K., Scarabel L., Qiu J., Zhu Q.-H., Yu Q., Timko M.P., Yamaguchi H., Merotto A., Qiu Y., Olsen K.M., Fan L., Ye C.-Y. (2022) Genomic insights into the evolution of Echinochloa species as weed and orphan crop. Nature Communications **13**:689.

Xiong Y., Yuan S., Xiong Y., Li L., Peng J., Zhang J., Fan X., Jiang C., Sha L., Wang Z., Peng X., Zhang Z., Yu Q., Lei X., Dong Z., Liu Y., Zhao J., Li G., Yang Z., Jia S., Li D., Sun M., Bai S., Liu J., Yang Y., Ma X. (2025) Analysis of allohexaploid wheatgrass genome reveals its Y haplome origin in Triticeae and high-altitude adaptation. Nature Communications **16**:3104.

Xu W.-Q., Ren C.-Q., Zhang X.-Y., Comes H.-P., Liu X.-H., Li Y.-G., Kettle C.J., Jalonen R., Gaisberger H., Ma Y.-Z., Qiu Y.-X. (2024) Genome sequences and population genomics reveal climatic adaptation and genomic divergence between two closely related sweetgum species. The Plant Journal **118**:1372–1387.

Yan Q., Wu F., Xu P., Sun Z., Li J., Gao L., Lu L., Chen D., Muktar M., Jones C., Yi X., Zhang J. (2021) The elephant grass (Cenchrus purpureus) genome provides insights into anthocyanidin accumulation and fast growth. Molecular Ecology Resources **21**:526–542.

Zea mays genome assembly Zm-B73-REFERENCE-NAM-5.0 NCBI [online] URL: https://www.ncbi.nlm.nih.gov/datasets/genome/GCF_902167145.1/ (accessed 21 September 2025).
Zhang G., Ge C., Xu P., Wang S., Cheng S., Han Y., Wang Y., Zhuang Y., Hou X., Yu T., Xu X., Deng S., Li Q., Yang Y., Yin X., Wang W., Liu W., Zheng C., Sun X., Wang Z., Ming R., Dong S., Ma J., Zhang X., Chen C. (2021) The reference genome of Miscanthus floridulus illuminates the evolution of Saccharinae. Nature Plants **7**:608–618.

Zhang J., Wu F., Yan Q., John U.P., Cao M., Xu P., Zhang Z., Ma T., Zong X., Li J., Liu R., Zhang Y., Zhao Y., Kanzana G., Lv Y., Nan Z., Spangenberg G., Wang Y. (2021) The genome of Cleistogenes songorica provides a blueprint for functional dissection of dimorphic flower differentiation and drought adaptability. Plant Biotechnology Journal **19**:532–547.

Zhang J., Zhang X., Tang H., Zhang Q., Hua X., Ma X., Zhu F., Jones T., Zhu X., Bowers J., Wai C.M., Zheng C., Shi Y., Chen S., Xu X., Yue J., Nelson D.R., Huang L., Li Z., Xu H., Zhou D., Wang Y., Hu W., Lin J., Deng Y., Pandey N., Mancini M., Zerpa D., Nguyen J.K., Wang L., Yu L., Xin Y., Ge L., Arro J., Han J.O., Chakrabarty S., Pushko M., Zhang W., Ma Y., Ma P., Lv M., Chen F., Zheng G., Xu J., Yang Z., Deng F., Chen X., Liao Z., Zhang X., Lin Z., Lin H., Yan H., Kuang Z., Zhong W., Liang P., Wang G., Yuan Y., Shi J., Hou J., Lin J., Jin J., Cao P., Shen Q., Jiang Q., Zhou P., Ma Y., Zhang X., Xu R., Liu J., Zhou Y., Jia H., Ma Q., Qi R., Zhang Z., Fang J., Fang H., Song J., Wang M., Dong G., Wang G., Chen Z., Ma T., Liu H., Dhungana S.R., Huss S.E., Yang X., Sharma A., Trujillo J.H., Martinez M.C., Hudson M., Riascos J.J., Schuler M., Chen L.-Q., Braun D.M., Li L., Yu Q., Wang J., Wang K., Schatz M.C., Heckerman D., Van Sluys M.-A., Souza G.M., Moore P.H., Sankoff D., VanBuren R., Paterson A.H., Nagai C., Ming R. (2018) Allele-defined genome of the autopolyploid sugarcane Saccharum spontaneum L. Nature Genetics **50**:1565–1573.

Zhang Y., Zhang G.-Q., Zhang D., Liu X.-D., Xu X.-Y., Sun W.-H., Yu X., Zhu X., Wang Z.-W., Zhao X., Zhong W.-Y., Chen H., Yin W.-L., Huang T., Niu S.-C., Liu Z.-J. (2021) Chromosome-scale assembly of the Dendrobium chrysotoxum genome enhances the understanding of orchid evolution. Horticulture Research **8**:183.

Zheng H., Wang B., Hua X., Gao R., Wang Y., Zhang Z., Zhang Y., Mei J., Huang Y., Huang Y., Lin H., Zhang X., Lin D., Lan S., Liu Z., Lu G., Wang Z., Ming R., Zhang J., Lin Z. (2023) A near-complete genome assembly of the allotetrapolyploid *Cenchrus fungigraminus* (JUJUNCAO) provides insights into its evolution and C4 photosynthesis. Plant Communications **4**:100633.

Zhu T., Wang L., Rimbert H., Rodriguez J.C., Deal K.R., De Oliveira R., Choulet F., Keeble-Gagnère G., Tibbits J., Rogers J., Eversole K., Appels R., Gu Y.Q., Mascher M., Dvorak J., Luo M.-C. (2021) Optical maps refine the bread wheat Triticum aestivum cv. Chinese Spring genome assembly. The Plant Journal **107**:303–314.
